# Immunosuppressants Rewire the Gut Microbiome-Alloimmune Axis Through Time-Dependent and Tissue-Specific Mechanisms

**DOI:** 10.1101/2025.01.02.631100

**Authors:** Long Wu, Allison Kensiski, Samuel J Gavzy, Yang Song, Hnin Wai Lwin, Michael France, Dejun Kong, Lushen Li, Ram Lakhan, Vikas Saxena, Wenji Piao, Marina W. Shirkey, Valeria R Mas, Brendan Lohmar, Yan Shu, Jonathan S Bromberg, Bing Ma

## Abstract

**Background:** Lifelong immunosuppressive therapy is required to prevent allograft rejection in organ transplantation. Current immunosuppressants effectively suppress adaptive and innate immune responses, but their broad, antigen-non-specific effects often result in severe off-target complications. It remains a significant unmet medical need in transplant medicine.

**Results:** In this study we investigated immunosuppressant effects of four major immunosuppressant classes, including tacrolimus, prednisone, mycophenolate mofetil (MMF), and fingolimod (FTY), on the gut microbiome, metabolic pathways, lymphoid architecture and lymphocyte trafficking after up to 30-day chronic exposure. Despite their distinct mechanisms of action and not designed to target the gut, all immunosuppressive drugs induced profound and time-dependent alterations in both intestine gene expression and gut microbiome composition. Progressive alterations from moderate early, drug-specific changes to a strikingly convergent microbial dysbiosis, marked by significant expansion of pathobionts of *Muribaculaceae*, occurred across all drug classes. Concurrently, all drugs uniformly induced significant suppression of mucosal immunity including B cell, immunoglobulin, and antigen recognition. Time-dependent changes in lymph node (LN) reorganization and cellular composition were also observed, marked by a progressive shift toward pro-inflammatory phenotypes in gut-draining mesenteric LNs and a gradual loss of tolerogenic architecture in peripheral LNs. Drug-specific metabolic alterations and distinct phases of intestinal transcriptional responses were also characterized. Notably, MMF and FTY demonstrated the most robust immunomodulatory properties, and were able to suppress alloantigen-induced inflammation through mediating regulatory T cell distribution and LN remodeling.

**Conclusions:** Together, these findings highlight the underappreciated complexity and temporal dynamics immunosuppressants effects, particularly their impact on the gut and compartmentalized regulation of alloimmune in lymphoid tissues. Understanding these relationships offers new opportunities for refining immunosuppressive strategies to reduce treatment-related off-target complications and improve long-term organ transplant outcomes.

## Introduction

Prevention of solid organ transplant rejection necessitates lifelong, multimodal immunosuppression to dampen both adaptive and innate alloreactive immunity. Despite improved short-term outcomes, long-term graft survival remains suboptimal due to chronic rejection and a range of adverse events such as metabolic complications, infections, neoplasia, and organ-specific toxicities ^1–8^. Current immunosuppressants target specific immune pathways but are limited by their antigen-non-specific nature, broadly suppressing the immune system and causing off-target effects ^9,10^. This dilemma between necessary immunosuppression and its imprecise, system-wide effects represents a critical challenge for improving long-term transplant outcomes.

A comprehensive understanding of the effects of immune suppressant drugs is essential for optimizing therapeutic regimens, minimizing adverse events, and preventing graft rejection. While most mechanistic studies on immunosuppressants have focused on their effects on T cells, B cells, and dendritic cells (DCs), little is known about their impact on secondary lymphoid organs (SLOs), particularly lymph nodes (LNs). As key regulators of local and systemic immunity, LNs direct immune responses toward tolerance or activation, with their function critically dependent on the architecture maintained by lymph node stromal cell (LNSC) networks, including fibroblastic reticular cells (FRCs), lymphatic endothelial cells (LECs), and blood endothelial cells (BECs). LNSCs orchestrate the positioning and interaction of lymphocytes with antigen-presenting cells (APCs) like DCs through the coordinated action of chemokines, cytokines, and stromal fibers ^11–13^. FRC-derived laminins are particularly important, as the laminin α4:α5 ratio (La4:La5) within the T cell cortex of the cortical ridge (CR) and around high endothelial venules (HEVs) modulates the balance between tolerance and immunity and influences global immune states ^14,15^. An increased La4:La5 promotes a pro-tolerogenic environment, whereas a decreased ratio fosters inflammation. LN structure and FRC function are often disrupted during transplantation due to ischemia-reperfusion injury (IRI) to the graft, leading to persistent “immunologic scarring”, a pathologic alteration of the LN cellular network ^16,17^. Further, regulatory T cells (Tregs) do not simply suppress immune responses through their presence alone. Their precise localization within LNs is critical for maintaining immune homeostasis. Tregs around HEVs limit the entry of pro-inflammatory effector T cells into the LNs. This gatekeeping function helps maintain a tolerogenic environment by preventing excessive accumulation of pro-inflammatory cells ^14^. Tregs in the CR suppress T cell priming and activation by APCs, preventing effector T cell activation and differentiation ^18^. A reduction of Tregs in these regions can result in enhanced T cell activation, promoting a pro-inflammatory state ^19–21^. Despite the importance of LN architecture and Treg localization in immune modulation, the effects of immunosuppressive drugs on these structural and functional elements remain poorly understood. Addressing this gap is critical for developing strategies to enhance long-term allograft survival.

Beyond their immunomodulatory effects, immunosuppressive drugs profoundly influence gut microbiome composition and function, which has been significantly implicated in alloimmunity and graft survival ^22–27^. Our group has demonstrated that gut microbiota modulates LN architecture by altering stromal fiber laminins ^28–30^. Our recent studies of tacrolimus and the mammalian target of rapamycin (mTOR) inhibitor rapamycin have revealed broad effects of these drugs on gut microbiome, luminal metabolic functions, intestinal transcriptome, as well as LN architecture ^31,32^, consistent with previous studies ^33,34^. This work suggests a crucial mechanistic link between gut microbiota and immune regulation. Other research has shown diverse mechanisms by which immunosuppressants affect host-microbe interactions. For example, methotrexate inhibits the conserved dihydrofolate reductase pathway in both host and gut bacteria, leading to broad microbiome alterations ^35^, while certain immunosuppressants disrupt the intestinal barrier, altering intestinal permeability ^36^. Notably, these host-microbe interactions are bidirectional, as gut microbiota can also alter intestinal function and drug bioavailability ^37,38^. For example, mycophenolate mofetil (MMF) is metabolized to inactive glucuronidated mycophenolic acid (MPAG) via hepatic glucuronidation, and luminal bacteria expressing β-glucuronidase can reactivate the metabolized form MPAG back to active MPA, leading to increased concentrations of active luminal drug and colonic inflammation ^39^. These complex interactions among immunosuppressants, gut microbiota, intestinal function, and immune cell populations underscore the need for a comprehensive understanding of their interrelationships to optimize therapeutic efficacy and mitigate unintended effects.

In this study, we investigated the understudied effects of immunosuppressants on LN architecture, lymphocyte and myeloid cell distribution, and the gut microbiome. We examined four key classes of clinical immunosuppressants: the calcineurin inhibitor tacrolimus (TAC), the glucocorticoid prednisone (PRED), the inosine monophosphate dehydrogenase (IMPDH) inhibitor MMF, and the sphingosine-1-phosphate (S1P) receptor antagonist fingolimod (FTY). Our findings revealed complex, time-dependent and tissue-specific patterns in drug responses and strikingly convergent gut microbial and intestinal responses. Despite their distinct mechanisms of action and not designed to target the gut, all drugs induced progressive alterations from moderate early changes to substantial alterations with prolonged treatment, converging to a shared dysbiotic state in the gut microbiome marked by significant expansion of pathobionts of *Muribaculaceae* across all drug classes. This was accompanied by significant metabolic alterations and distinct phases of intestinal transcriptional responses. Time-dependent changes were also observed in LNs. The mesenteric LNs (mLNs) developed progressively increased pro-inflammatory states with extended immunosuppressive use, while peripheral LNs (pLNs) showed strong early pro-tolerogenic effects that became less pronounced over time. While this compartmentalized immune regulation was broadly consistent across all drugs studied, MMF and FTY demonstrated particularly robust immunomodulatory effects. These two agents effectively suppressed alloantigen-induced pro-inflammatory changes, through mediating Treg redistribution and LN remodeling. Overall these findings revealed the underappreciated complexity and dynamics of immunosuppressant effects, suggesting a mechanistic link between immunosuppressant-induced gut dysbiosis, compartmentalized immune regulation in peripheral and mesenteric lymphoid tissues, and immunosuppressant off-target complications.

## Results

### Time-dependent effects of immunosuppressants on gut microbiome

We treated groups of C57BL/6 mice daily with four immunosuppressants (TAC, PRED, MMF and FTY), respectively, using untreated mice as controls. We analyzed the gut microbiome at early (3 days), intermediate (7 days), and late (30 days) time points. We performed whole community metagenomic sequencing on colonic intraluminal stool samples and utilized the comprehensive mouse gut metagenome catalogue (CMGM) ^40^ (**Supplemental Table 1A**) for taxonomic profiling and Human Microbiome Project Unified Metabolic Analysis Network (HUMAnN3) ^41^ (**Supplemental Table 1B**) to analyze microbial metabolic pathways.

To quantify immunosuppressant drug effects on the gut microbiome, we used Bray-Curtis distance analyses to measure changes in both taxonomic composition and metabolic pathways (**Figure 1A, B**). We compared each treatment group to untreated controls (treatment-to-control) to determine deviations from baseline and analyzed differences between different treatment groups (treatment-to-treatment). In treatment-to-control pairwise comparisons, the effects were variable during early treatment by day 3 and 7. However, by day 30, all treatment groups showed profound and significant alterations in their gut microbiome profiles, suggesting a cumulative drug effect with prolonged treatment. The pattern was evident in both taxonomic compositions and functional pathways, with the most substantial changes emerging with 30-day treatment. Drug-to-drug pairwise comparison also revealed distinct time-dependent drug effects on gut microbiome. The differences between treatment groups increased from day 3 to day 7, indicating drug-specific early effects on microbial communities. However, by day 30, these inter-drug differences had substantially decreased, suggesting that despite their distinct mechanisms of action, prolonged immunosuppression lead to convergent effects on the gut microbiome. These findings were further supported by ordination analyses. Principal Component Analysis (PCA) revealed clear clustering of 30-day samples, distinct from earlier time points, regardless of the specific immunosuppressant used (**Figure 1C**). Canonical Correspondence Analysis (CCA) confirmed these observations, demonstrating significant correlations between treatment duration and microbiome composition, with maximal separation occurring at 30 days (**Supplemental Figures 1A and 1B**).

**Figure 1:**
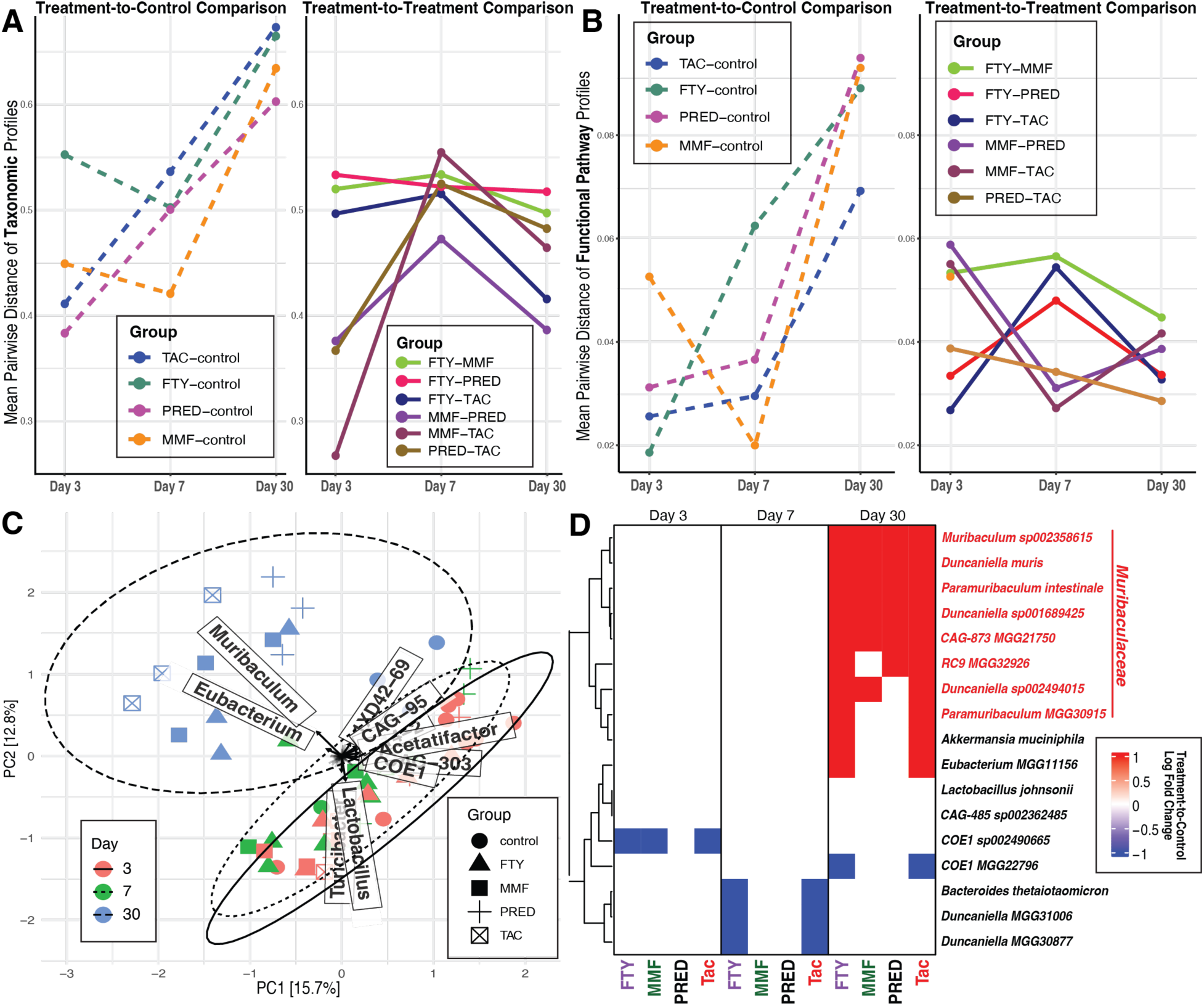
Impact of immunosuppressant treatment on gut microbiome composition and function. Pairwise Bray-Curtis distance analysis comparing drug-treated groups to untreated controls and between different treatment groups to measure changes in **A)** taxonomic composition, and **B)** functional pathway abundance. Distances represent mean values per group. Untreated mice as control group. **C**) Principal Components Analysis of gut microbiome composition based on Bray-Curtis distance. Robust centered log ratio transformation applied on count compositional data. **D)** Differentially abundant taxonomic groups identified by pairwise comparisons between treatment groups and controls. Changes in abundance are expressed as log2 fold changes relative to control group.

The temporal convergence of microbiome alterations across different immunosuppressant drugs revealed shared downstream influences of these drugs on host-microbe interactions, despite their diverse initial effects. To identify microbial signatures associated with immunosuppression, we compared the abundance of bacterial species between drug-treated and control groups across multiple timepoints to reveal treatment-specific and temporal changes in microbial populations (**Figure 1D**). No individual taxonomic group showed consistent alterations across all timepoints in any treatment group. Early (day 3) and intermediate (day 7) time points revealed sporadic alterations in various bacterial taxa, but these changes were transient. By day 30, all treatment groups, despite their distinct mechanisms of action, converged on a common set of bacterial taxa that were not significant at the earlier time points. These taxa belonged exclusively to the family of *Muribaculaceae* (formerly known as *S24-7*), including *Duncaniella*, *Paramuribaculum intestinale*, and *CAG-873*. This family is known for its metabolic versatility and adaptability ^42–44^, and is primarily a gut commensal in mice but can act as pathobionts under dysbiotic conditions, with recent evidence demonstrating their causative role in insulin-dependent diabetes in mouse models ^45^. Their enrichment indicates a shift toward a dysbiotic state in which the microbial balance is skewed toward species with pathobiont potentials with altered metabolic capabilities. This progression from early drug-specific changes to a convergent dysbiotic state with prolonged treatment underlines the profound and progressive impact of chronic immunosuppressant use on the gut microbiome.

### Conserved intestinal responses to immunosuppressants revealed induction of epithelial stress and suppression of lymphoid signatures

To understand host gut responses to drug treatment, we analyzed the transcriptome of small intestine at days 7 and 30, as these time points corresponded to significant alterations in the gut microbiome during treatments. Differentially expressed genes (DEGs) were identified by comparing each drug treatment group to the no-treatment controls (**Supplemental Table 2A**). DEGs analysis revealed that 30-day treatments induced 1.3 to 2 times more DEGs than 7-day treatments (**Figure 2A**), indicating that prolonged immunosuppression triggered more extensive transcriptional changes. At day 7, FTY induced the largest number of DEGs. By day 30, both FTY and PRED showed stronger effects in both upregulation and downregulation of DEGs. In contrast, MMF showed the least impact, particularly in upregulated DEGs. TAC induced the largest portion of upregulated DEGs with twice as many upregulated DEGs as downregulated ones, compared to other drugs that showed either greater downregulation or balanced regulation. Overall, prolonged immunosuppressive treatment drove broader and more extensive transcriptional changes in intestinal tissue.

**Figure 2:**
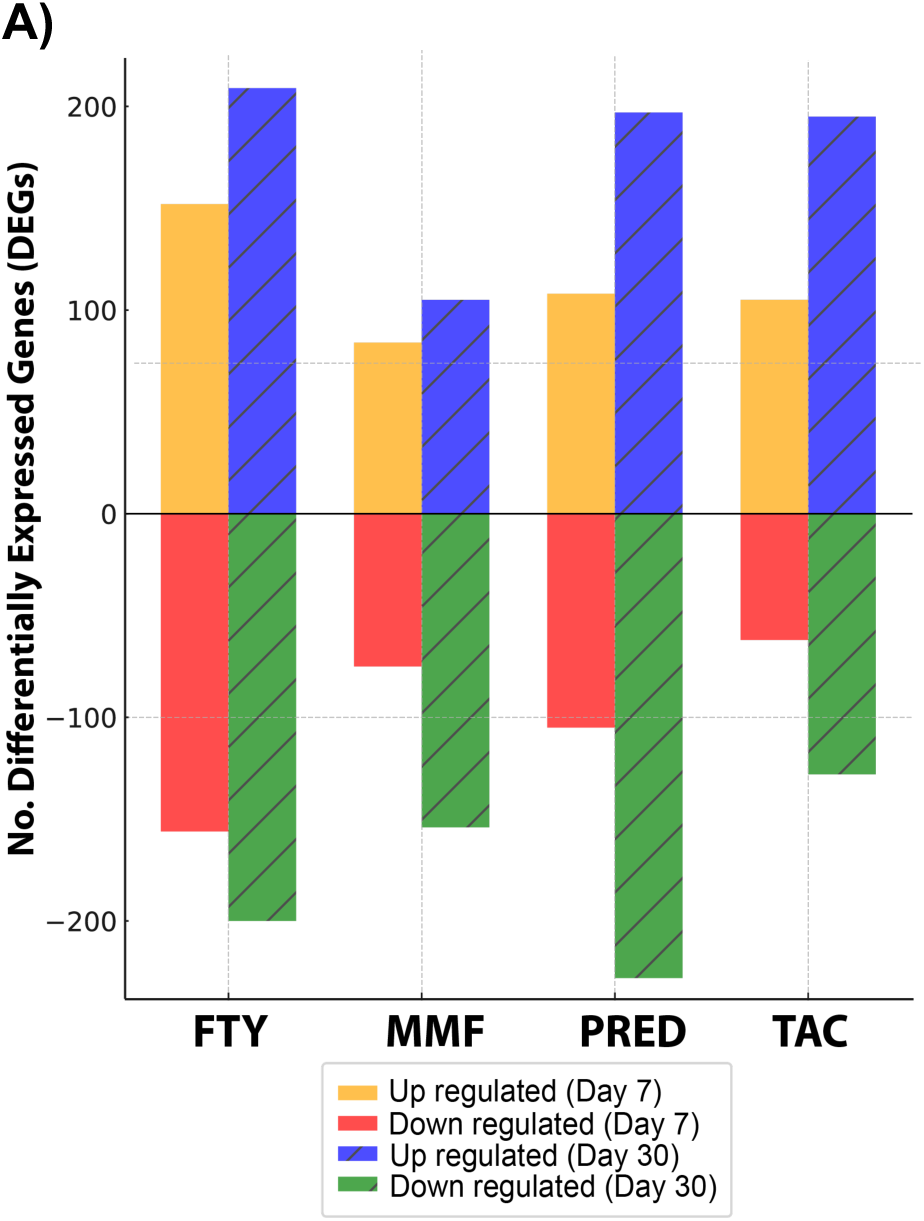
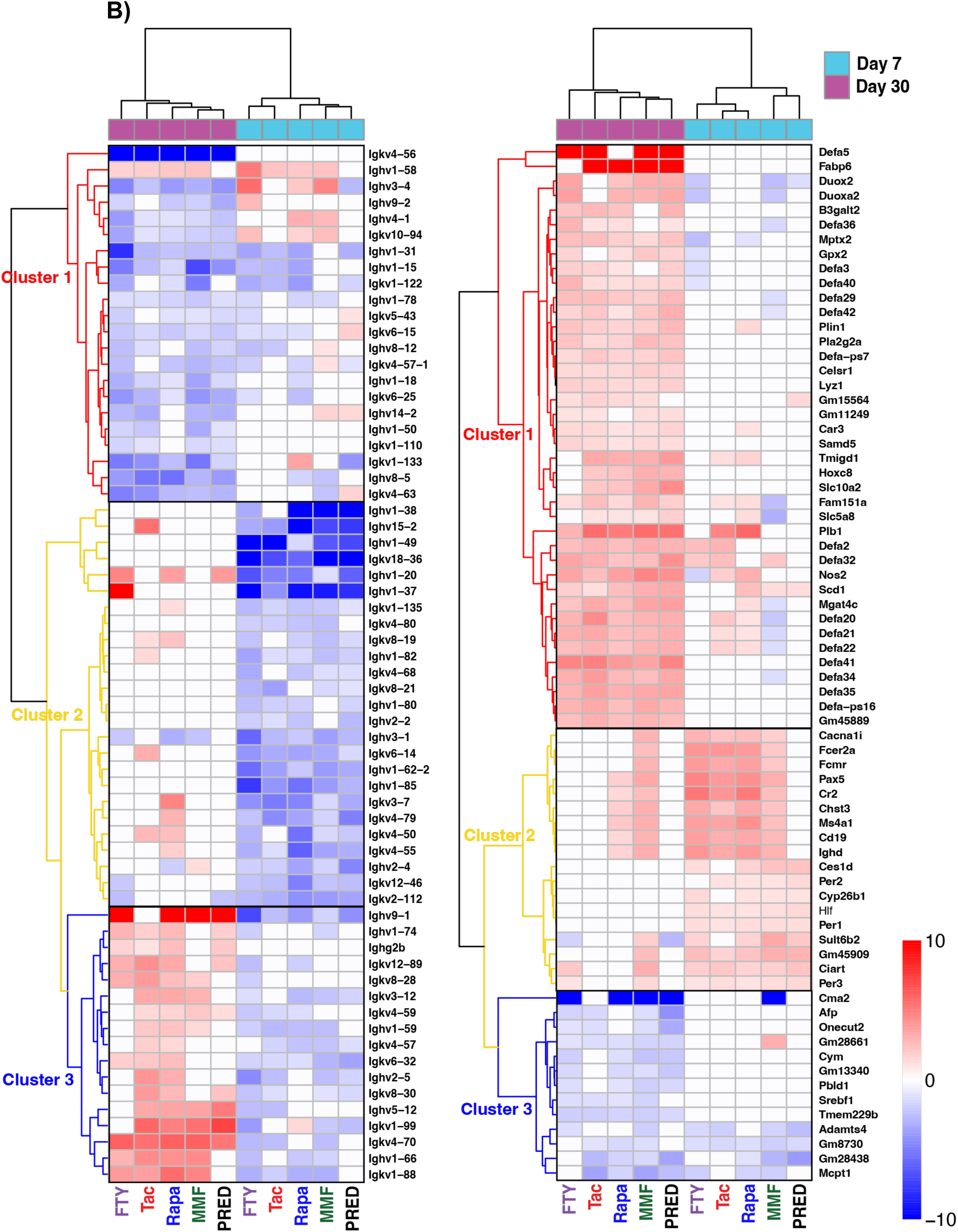
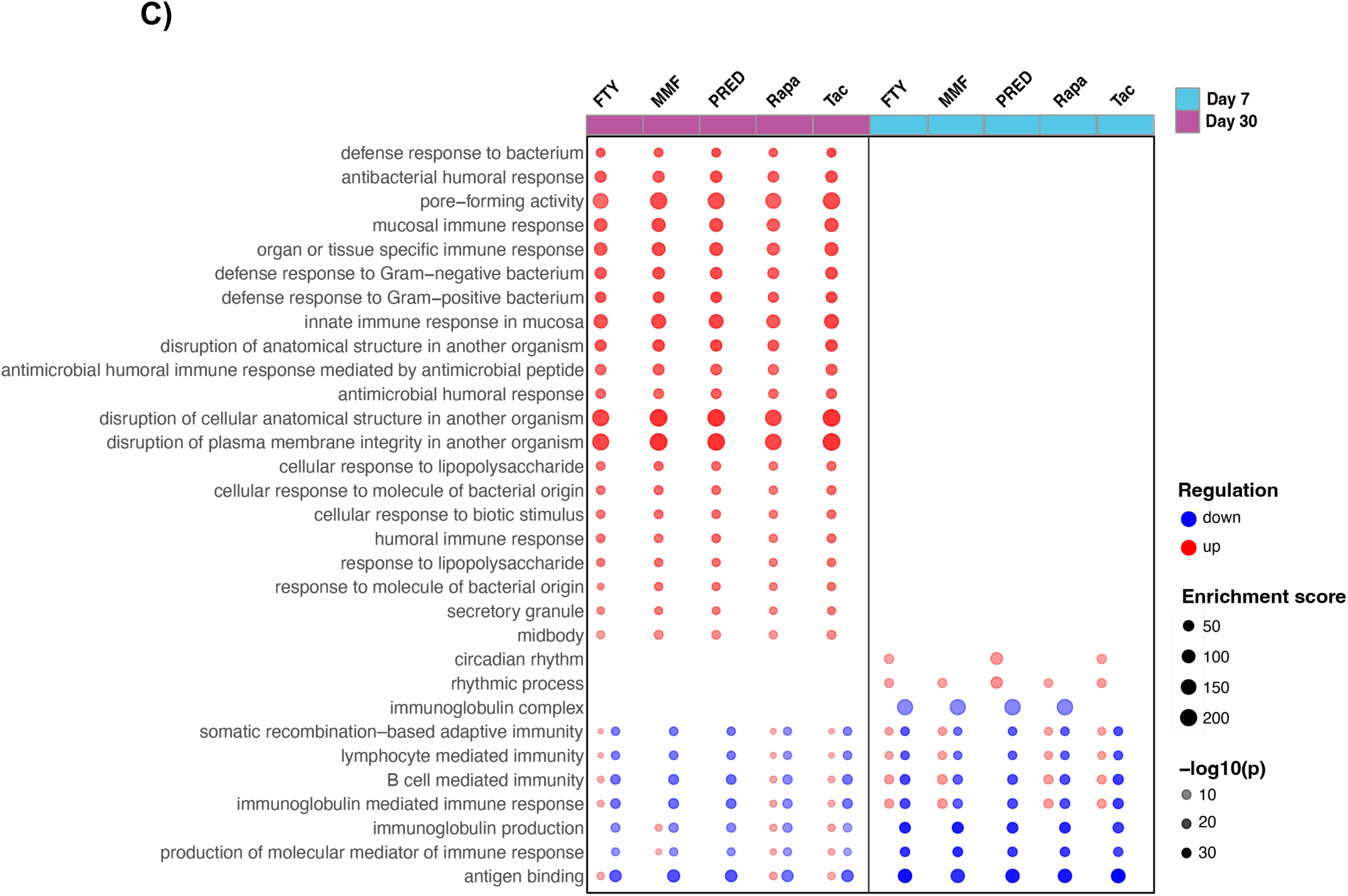
Immunosuppressants temporally shift intestinal immune responses. **A**) Bar plots showing the number of differentially expressed genes (DEGs) after 7 and 30 days of treatment compared to the control group. Yellow and blue bars indicate the number of upregulated genes, while red and green bars represent the number of downregulated genes. **B**) Heatmap DEGs in intestinal tissue following treatment with five immunosuppressant drugs at days 7 and 30. Transcriptomic data from rapamycin-treated mice from our previously study ^31^ integrated in current analysis. Genes defined as differentially expressed in at least four of the five drug treatment groups at either time point. Hierarchical clustering performed on both genes (rows) and samples (columns) using Euclidean distance and ward linkage. Color scale indicates normalized Z-scores of expression values, with red denoting upregulation and blue denoting downregulation relative to no-treatment controls. **C**) Functional enrichment analysis of conserved intestinal transcriptional responses to immunosuppressant treatments. Dot plot showing over-represented Gene Ontology (GO) biological process pathway based on DEGs at days 7 and 30. EnrichGO pathways containing more than three genes were included in this plot. Dot size denotes the normalized enrichment score. Blue downregulated and red upregulated pathways; color shaded based on the –log₁₀(p-value) of enrichment. **Abbr**: TAC: tacrolimus, MMF: mycophenolate mofetil, PRED: prednisone, FTY: fingolimod, RAPA: rapamycin

We next analyzed DEGs common to all immunosuppressant treatments to identify shared drug-induced effects (**Supplemental Table 2B-D**). To enable a comprehensive comparison across different immunosuppressant classes, we integrated transcriptomic data from rapamycin-treated mice from our previously study ^31^ into the current analysis (**Figure 2B**). Conserved DEGs were defined as genes consistently differentially expressed by at least four drugs at either time points. Clustering analysis of these conserved DEGs at days 7 and 30 revealed distinct temporal expression patterns, particularly affecting genes involved in mucosal immunity, epithelial homeostasis, metabolism, and circadian regulation (**Figure 2C**).

Specifically, immunoglobulin heavy variable (Ighv) and kappa light chain variable (Igkv) region genes displayed broad yet temporally distinct responses under immunosuppressant treatments. Cluster 1 DEGs (**Figure 2B** left panel) includes genes mainly suppressed at day 30, including classical Ighv and Igkv segments typically associated with naive and T-dependent germinal center-derived B cells. This downregulation indicates a significant, prolonged suppression of *de novo* B cell activation and diversification, especially affecting clones that require antigen-specific, T cell-dependent signals. Cluster 2 DEGs were mostly suppressed at day 7 only. These genes, which include variable region segments commonly expressed by naive or early responding B cells, suggest an early transcriptional quiescence. In contrast, Cluster 3 genes were initially suppressed at day 7 but later upregulated at day 30. The activation of the Ighg2b gene suggests that class-switch recombination has occurred, reflecting the selective survival and expansion of mucosal or microbiota-tolerant B cell subsets capable of adapting to the chronically immunosuppressed gut. Overall, these results indicate a predominantly suppressed B cell transcriptional program, with limited recovery by day 30.

Further immunohistochemical (IHC) assessments of intestinal conventional dendritic cells (CD11c+ cDCs) and regulatory T cells (Foxp3+ Tregs) revealed day 7 was the critical transitional point for all immunosuppressant drugs (**Supplemental Figure 2A-C**). A significant reduction in intestinal cDC populations at day 7 was observed for all drugs, with no significant alterations observed at days 3 or 30. Intestinal Tregs similarly had the most significant decreases at day 7, while returning to baseline levels by day 30. Collectively, these results indicated that at day 7, B cells experienced diminished antigenic stimulation, reduced survival and differentiation signals, and suppressed transcription of immunoglobulin variable region genes. By day 30, limited restoration of antigen delivery, B and T cell costimulatory signals, and epithelial tolerance to baseline levels associated with partial resumption of recombined immunoglobulin gene transcription, especially among surviving tissue-resident B cells. This partially restored yet still broadly suppressed state may represent a compensatory host response, favoring class-switched memory or microbiota-reactive B cell clones to maintain mucosal homeostasis under ongoing immunosuppressive therapy.

Epithelial DEGs showed three distinct temporal expression clusters, reflecting the mucosal defense and host compensatory responses to immunosuppressant treatment. Cluster 1 DEGs were significantly activated by day 30, indicative of robust mucosal defense mechanisms and adaptive metabolic programs. Outstandingly, this cluster included key antimicrobial effectors, including α-defensins (Defa-ps16, Defa-ps7, Defa2, Defa20-22, Defa29, Defa3, Defa32, Defa34-36, Defa40-42, Defa5), lysozyme (Lyz1), and secretory phospholipase A2 group IIA (Pla2g2a), which are canonical products of Paneth cells ^46^. This transcriptional profile is consistent with a Paneth cell-driven antimicrobial response, suggesting that activation of innate epithelial defense mechanisms was possibly triggered by pathobiont expansion and microbiota perturbation following prolonged immunosuppressant exposure. Further, oxidase genes (Duox2, Duoxa2) were elevated, supporting reactive oxygen species (ROS) production for microbial control. Moreover, upregulation of inducible nitric oxide synthase (iNOS, Nos2) and glutathione peroxidase (Gpx2) suggested enhanced epithelial stress responses. These adaptive responses likely represented host mechanisms to control potential microbiota perturbations. Cluster 2 genes, mainly upregulated at day 7, prominently included key circadian regulators (Ciart, Per1, Per2, Per3, Hlf), suggesting an early host-driven resetting of circadian rhythms in response to drug-induced gut perturbation. Further, genes involved in xenobiotic metabolism and detoxification pathways (Cyp26b1, Ces1d, Sult6b2) and early immune interaction genes (Cd19, Fcer2a, Pax5) were also significantly induced. Together these genes represented an early epithelial, metabolic and immune adaptation to drug disturbance. Cluster 3 genes were downregulated at day 30, including key genes involved in epithelial differentiation and renewal (Onecut2, Tmem229b), lipid biosynthesis (Srebf1), extracellular matrix remodeling (Chst3, Adamts4), and mast cell proteases (Mcpt1, Cma2). The suppression of these functions reflected a compromised capacity for epithelial regeneration and reduced structural adaptability after prolonged treatment. Overall, these data showed a profound off-target influence of the immunosuppressant drugs on the intestine, and importantly, a dynamic mucosal adaptation to immunosuppressive induction, transitioning from initial circadian and xenobiotic metabolism, progressing toward robust innate mucosal defense programs and suppression of epithelial renewal and structural remodeling processes.

### Drug-specific alterations in intestinal expression and gut luminal metabolic activity

Beyond shared mechanisms, we next examined drug-specific transcriptional programs to identify unique intestinal responses elicited by individual immunosuppressant agents. UpSet plots were used to analyze DEGs unique to or shared between days 7 and 30, as well as across different treatment groups (**Supplemental Figure 3A-D**). Drug effects were more pronounced at day 30, with 1.6-3.3 times more down-regulated DEGs and 1.4-2.2 times more upregulated DEGs compared to day 7. Among the treatments, TAC showed the most pronounced effect at day 30, while FTY had the least impact. Further, minimal overlap in DEGs was observed between timepoints and across different treatments, indicating transient effects at day 7. For instance, following PRED treatment, only 31 genes (15 upregulated, 16 downregulated) were consistently modulated at both timepoints, while 255 genes were uniquely regulated at day 7 (81 upregulated, 78 downregulated) and 371 at day 30 (174 upregulated, 197 downregulated). This pattern held true for all four drug treatments. Further comparison between treatment groups revealed distinct patterns of gene expression (**Supplemental Figure 4A-B**). The overlap in DEGs was limited, with less than 20% between any two drugs and less than 5% across the four drugs, underscoring highly drug-specific mechanisms of action on intestine. Together, these findings demonstrated that immunosuppressants induced time-dependent intestinal responses, with most pronounced drug-specific changes following prolonged treatment.

Gene ontology enrichment analyses of DEGs revealed drug-specific patterns (**Supplemental Figure 4C**). At day 7, FTY showed the strongest suppressive effect on mucosal and humoral immunity pathways, which transitioned to moderate activation of immunoglobulin production by day 30. MMF shifted from early activation on day 7 to broad suppression of innate immune pathways at day 30. TAC showed moderate increases in lymphocyte and immunoglobulin pathways at day 7, progressing to stronger activation of both innate immunity and antibody production at day 30. PRED maintained relatively modest effects throughout the treatment period, though the specific pathways affected differed between the two timepoints. Together, these data demonstrated that immunosuppressant drugs exerted both shared and drug-specific effects on the intestine, profoundly impacting epithelial function as well as innate and adaptive immune pathways, contributing to the off-target effects of immunosuppressive therapies.

Gut luminal metabolic activity of small intestine was further analyzed at day 7, as this time point represented a critical transitional phase. Untargeted metabolomic analysis identified 264 distinct metabolites in intraluminal stool samples (**Supplemental Table 3A-B**). Partial Least Squares Discriminant Analysis (PLS-DA) demonstrated separations of metabolic profiles among the different treatment groups (**Supplemental Figure 5A**), and the top contributing metabolites in each group are shown in **Supplemental Figure 5B**. Hierarchical clustering analysis further confirmed these drug-specific patterns, demonstrating that each immunosuppressant induced unique changes in metabolite abundance (**Supplemental Figure 5C**). Pairwise comparison of individual drugs to no-treatment controls, combined with enrichment analyses, revealed both shared and drug-specific effects on metabolic activities (**Table 1**, **Supplemental Table 3C-F**).

**Table 1.** Differential metabolite abundance in gut lumen after 7 days of immunosuppressant treatment *.

|  | TAC | PRED | MMF | FTY |
| --- | --- | --- | --- | --- |
| ↑ | **31**<br>• Amino acids & derivatives<br>• Pyrimidine metabolism<br>• Sugars and sugar phosphates | **4** | **4** | **14**<br>• Purine metabolism<br>• Arginine metabolism |
| ↓ | **28**<br>• Polyamines & derivatives<br>• SCFAs, TMA | **95**<br>• Amino acids & derivatives<br>• SCFAs, TMA<br>• Pyrimidine metabolism<br>• Polyamines & derivatives | **37**<br>• Glyoxylate and Dicarboxylate<br>• Branched-chain amino acids & derivatives<br>• Purine metabolism<br>• Neurotransmitters | **22**<br>• Pyrimidine metabolism<br>• SCFAs |
| *subtotal* | **59** | **99** | **41** | **36** |

\* Gut luminal metabolites with significant changes (fold change >2, p<0.05) after 7 days of treatment with tacrolimus (TAC), prednisone (PRED), mycophenolate mofetil (MMF), or Fingolimod (FTY) compared to untreated controls. Numbers in bold indicate total metabolites significantly increased (↑) or decreased (↓) for each drug. Key metabolic pathways affected are listed for each drug. SCFAs: short-chain fatty acids; TMA: trimethylamine. Data derived from untargeted metabolomics analysis using capillary electrophoresis-mass spectrometry.

TAC treatment induced similar numbers of significantly increased and decreased metabolites (**Supplemental Figure 6A, Supplemental Table 3F**). Metabolites elevated by TAC included amino acids and derivatives, suggesting altered protein catabolism and amino acid utilization; pyrimidine metabolites indicative of increased nucleotide turnover, and sugars/sugar phosphates reflecting enhanced glycolytic activity and central carbon metabolism. Conversely, TAC significantly reduced polyamines, biogenic amines, short-chain fatty acids (SCFAs), betaine, trimethylamine (TMA), and various organic acids. In contrast, PRED, MMF, and FTY primarily induced suppressive metabolic effects, each with distinct scope and profiles. PRED caused the broadest suppression, reducing three to four times more metabolites than it increased (**Supplemental Figure 6B, Supplemental Table 3D**), including amino acid derivatives, SCFAs, TMA, pyrimidine intermediates, and polyamines, consistent with its widespread inhibitory effects on cellular metabolism ^47–49^. MMF showed a similarly suppressive but more targeted profile, predominantly affecting glyoxylate/dicarboxylate metabolism, branched-chain and aromatic amino acids, and purine pathways (**Supplemental Figure 6C, Supplemental Table 3E**). MMF increased adenosine while reducing uric acid, consistent with its mechanism as an inosine monophosphate dehydrogenase (IMPDH) inhibitor ^50,51^, directly linking metabolic changes to its therapeutic effects. FTY induced a relatively milder suppressive profile (**Supplemental Figure 6D, Supplemental Table 3C**), with increased nucleosides and reduced nucleotides and SCFAs. Notably, decreased arginine and elevated N^1^,N^8^-diacetylspermidine and spermidine pointed to enhanced polyamine biosynthesis. Collectively, these findings underscore shared and distinct metabolic alterations in the gut lumen that may contribute to both therapeutic efficacy and off-target effects.

### Tissue-specific and time-dependent effects of immunosuppressants on mLN and pLN organization

We next investigated how immunosuppressants affected LN organization in both mucosal and systemic compartments by examining mLN and pLN, the primary draining sites for mucosal and systemic immunity, respectively. IHC provides spatial resolution, enabling localized assessments of Tregs within special areas such as LN HEVs and CRs, whereas flow cytometry offers a quantitative measurement of overall Treg populations. These methods complemented each other, measuring different aspects of immune cell dynamics and distribution. Using flow cytometry and IHC, we analyzed changes in the numbers and distribution of immune cell populations (B220+ B cells, CD4+ T cells, CD8+ T cells, Foxp3+ Tregs) and structural features (La4:La5 in LNs). We particularly assessed the changes to La4:La5 and distribution of Tregs in the CR and around the HEVs of LNs to understand how immunosuppressants modulated cellular and structural aspects of immune organization crucial for immune regulation.

In mLNs, flow cytometry revealed that B220+ B cells, CD4+ T cells, CD8+ T cells and Foxp3+ Tregs were unaffected by TAC or MMF treatment at days 3, 7, or 30; in contrast, PRED treatment significantly reduced CD4+ T cells and B220+ B cells by day 30, with no significant changes at earlier time points (**Figure 3A, C**). FTY treatment increased Foxp3+ Tregs by day 30, with no changes at days 3 or 7 (**Figure 3B, C**). Overall, while CD8+ T cell populations did not significantly change across conditions (data not shown), other populations showed treatment-specific changes that were only evident after prolonged treatment. IHC revealed two groups of change patterns. On the one hand, TAC and PRED showed no significant changes in Treg distribution at any time point (**Figure 3D, E, H, I, Supplemental Figure 6A, B**), but reduced La4:La5 around the CR after 30 days (**Figure 3F, H, Supplemental Figure 6C**). TAC additionally decreased this ratio in the HEV on day 7 (**Figure 3G, I, Supplemental Figure 6D**). On the other hand, MMF and FTY showed similar patterns with an increased Treg distribution in the CR by day 7, followed by a decrease by day 30 (**Figure 3D, H, Supplemental Figure 6A**) and a decreased La4:La5 in both CR and HEV (**Figure 3F-I, Supplemental Figure 6C, D**). Early responses at day 3 were mixed, suggesting initial variable effects converged to a stable state with prolonged drug treatment. Overall, despite their distinct mechanisms of action, all four immunosuppressant drugs ultimately induced a more pro-inflammatory environment in mLN by day 30, characterized by reduced Treg presence and/or decreased La4:La5. This temporal progression suggested that prolonged immunosuppression may paradoxically create localized pro-inflammatory conditions in mucosal-associated lymphoid tissues.

**Figure 3:**
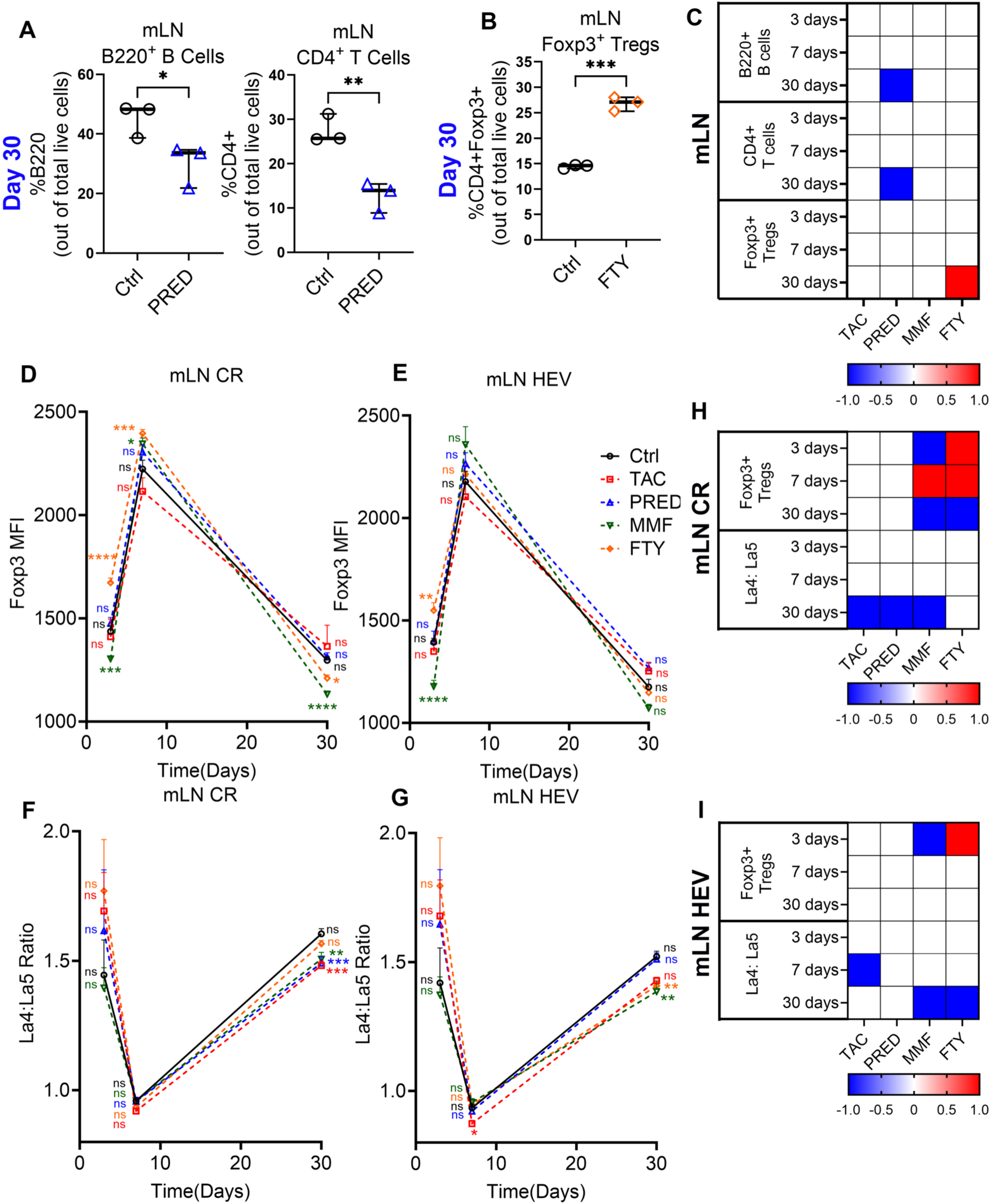
Effects of TAC, PRED, MMF, and FTY on mLN cell content, cell distribution, and structure. Flow cytometry analysis of **A)** B220+ B cells and CD4+ T cells on day 30 after PRED treatment and **B)** Foxp3+ Tregs on day 30 after FTY treatment in mLNs. **C)** Heatmaps depict changes in expression with TAC, PRED, MMF, and FTY relative to controls in mLNs after 3, 7, and 30 days of treatment. IHC of **D)** CR and **E)** HEV Foxp3+ Tregs on days 3, 7, and 30 in mLNs. IHC of **F)** CR and **G)** HEV La4: La5 on days 3, 7, and 30 in mLNs. Barplot showing detailed plots at each timepoint are in **Supplemental Figure 6**. Heatmaps depict changes in expression with TAC, PRED, MMF, and FTY relative to controls in **H)** CR and **I)** around HEV in mLNs. 3 mice/group. 2-3 LNs/mouse, 2-3 sections/LN, 7-30 fields/tissue. One-way ANOVA. * p < 0.05; ** p < 0.01, *** p < 0.001, **** p < 0.0001.

In pLNs, distinct temporal patterns of immune modulation were also observed. Using flow cytometry, TAC and PRED induced minimal effects on cellular composition (**Figure 4D**), and MMF selectively reduced CD4+ T cells by day 30 (**Figure 4A, D**). FTY showed the most dynamic changes, increasing B220+ B cells while decreasing CD4+ T cells at day 7 (**Figure 4B, D**), followed by elevated Foxp3+ Tregs and decreased CD4+ T cells by day 30 (**Figure 4C, D**). IHC analysis revealed complementary changes in cellular distribution and stromal architecture. TAC showed transient effects, characterized by increased Treg distribution in the CR at day 7 (**Figure 4E, I, Supplemental Figure 7A**) without affecting laminin ratios (**Figure 4G-J, Supplemental Figure 7C, D**). PRED demonstrated only early structural changes, marked by increased La4:La5 at day 3 **(Figure 4E-J, Supplemental Figure 7A-D)**. MMF showed the most consistent pro-tolerogenic profile, maintaining increased Treg distribution throughout the treatment period **(Figure 4E, F, I, J, Supplemental Figure 7A, B)** and a higher La4:La5 at day 3 **(Figure 4G-J, Supplemental Figure 7C, D)**. FTY showed early pro-tolerogenic effects with increased Treg distribution and La4:La5 at day 3, but not at later time points **(Figure 4E-J, Supplemental Figure 7A-D)**. Together, these findings demonstrated that all four immunosuppressants, particularly MMF and FTY, predominantly promoted a pro-tolerogenic state in pLNs, characterized by increased Tregs and/or elevated laminin α4:α5 ratios particularly during early treatment phases. This contrasted with the pro-inflammatory environment observed in mLNs, suggesting differential regulation of immune organization between loco-regional and systemic lymphoid tissue compartments.

**Figure 4:**
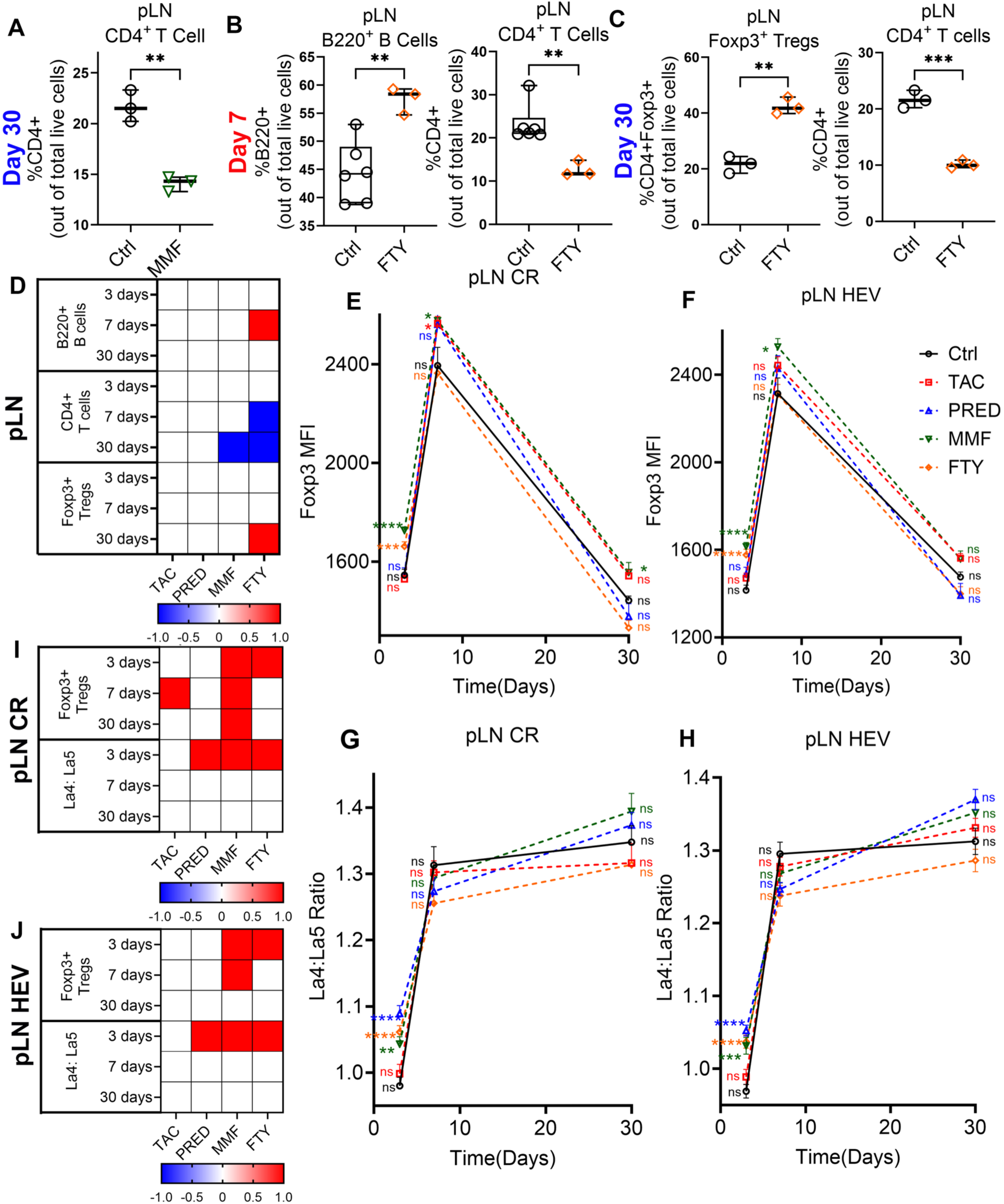
Effects of TAC, PRED, MMF, and FTY on pLN cell content, cell distribution, and structure. Flow cytometry analysis of **A)** CD4+ T cells on day 30 after MMF treatment, and **B)** B220+ B cells and CD4+ T cells on day 7 and **C)** CD4+ T cells and Foxp3+ Tregs after FTY treatment in pLNs. **D)** Heatmaps depict changes in expression with TAC, PRED, MMF, and FTY relative to controls in pLNs after 3, 7, and 30 days of treatment. IHC of **E)** CR and **F)** HEV Foxp3+ Tregs on days 3, 7, and 30 in pLNs. IHC of **G)** CR and **H)** La4: La5 on days 3, 7, and 30 in pLNs. Barplot showing detailed plots at each timepoint are in **Supplemental Figure 7**. Heatmaps depict changes in expression with TAC, PRED, MMF, and FTY relative to controls in **I)** CR and **J)** around HEV in pLNs. 3 mice/group. 2-3 LNs/mouse, 2-3 sections/LN, 7-30 fields/tissue. One-way ANOVA. * p < 0.05; ** p < 0.01, *** p < 0.001, **** p < 0.0001.

### MMF and FTY counteract alloantigen-induced pro-inflammatory changes through dual regulation of Tregs and LN architecture

Given the pronounced effects of MMF and FTY on LN organization under homeostatic conditions (**Figure 3H, I, 4I, J**), we next utilized a mouse model with allogeneic stimulation (Allo) to assess their impact on transplant-related alloimmune responses, mimicking the immunologic stress of solid organ transplantation. Mice were injected with fully allogeneic splenocytes (10^7^ cells intravenously) followed by MMF or FTY administration for 3 days. Compared to the no treatment controls, Allo alone did not affect Treg distribution in mLNs or pLNs (**Figure 5A, C**). However, when combined with either drug, significant increases in Tregs were observed. Compared to Allo alone, MMF increased Tregs around mLN HEV, and FTY increased Tregs in both mLN HEV and pLN HEV and CR (**Figure 5A, C, E, F**). Allo alone decreased La4:La5 in both pLN and mLN (**Figure 5B, D**), indicating a pro-inflammatory shift caused by alloantigen-induced immune responses, consistent with our previous findings ^15^. When combined with MMF or FTY, both drugs prevented this decrease, maintaining laminin α4:α5 ratios comparable to untreated controls **(Figure 5B, D, E, F)**. These findings demonstrated that MMF and FTY inhibited alloantigen-induced pro-inflammatory shifts in LN through two complementary mechanisms: increasing Tregs and preserving tolerogenic laminin stromal architecture. This dual action suggested that these drugs may be particularly effective at promoting a tolerogenic environment during allogeneic immune challenges.

**Figure 5.**
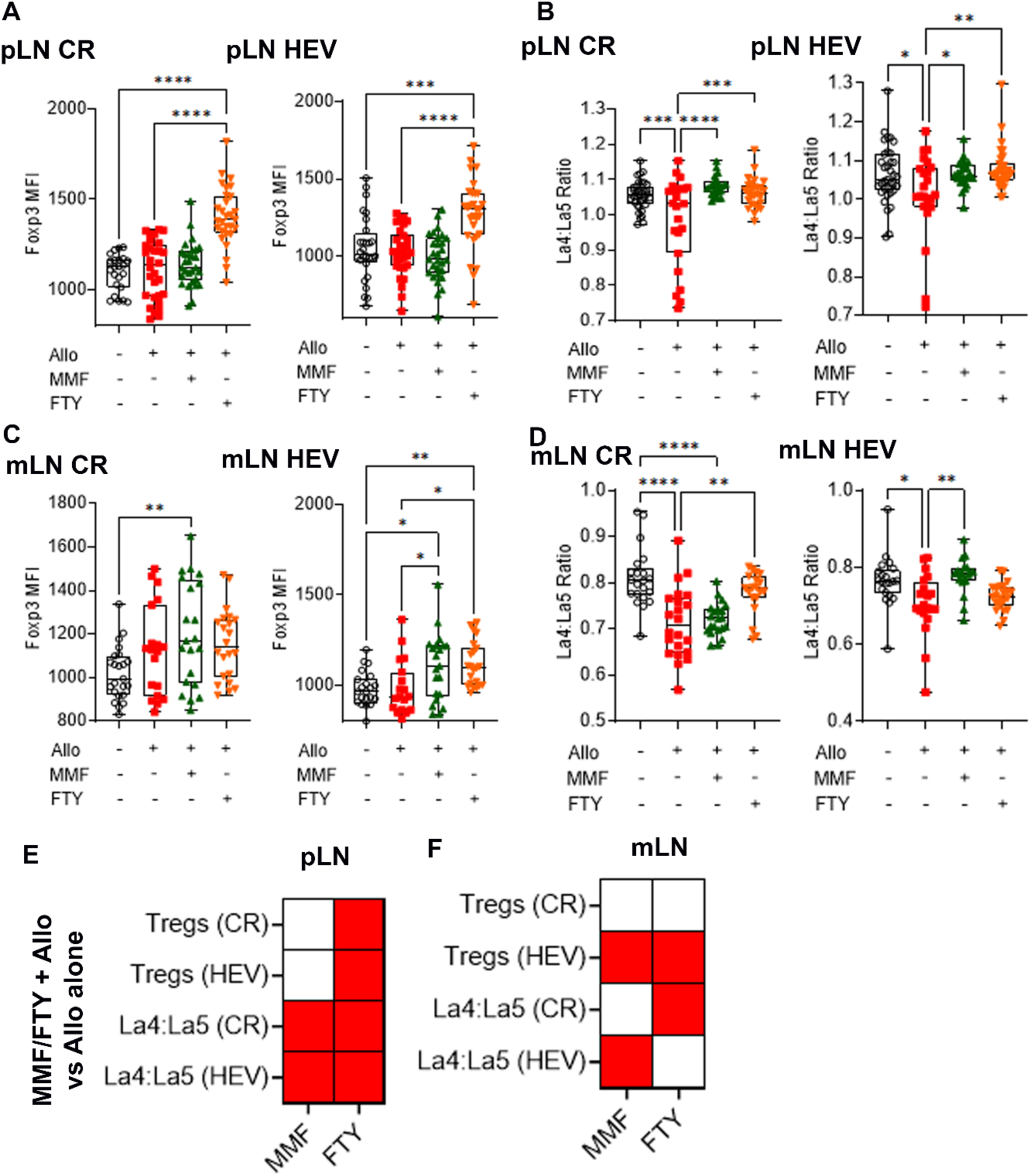
MMF and FTY induce a rapid anti-inflammatory response during allogeneic stimulation. IHC for Foxp3+ Tregs in the **A**) pLN and **C**) mLN. La4:La5 in **B**) pLN and **D**) mLN. Heatmap summarizes significant changes in expression comparing allogeneic stimulation plus MMF or FTY to allogeneic stimulation alone in **E**) pLNs and **F**) mLNs. 3 mice/group. 2-3 LNs/mouse, 2-3 sections/LN, 7-30 fields/tissue. One-way ANOVA: * p < 0.05, ** p < 0.01, *** p < 0.001, **** p < 0.0001.

### MMF and FTY effects are mediated through FRC-derived laminins α4 and α5

Given the significant impact of MMF and FTY on LN architecture, particularly through modulation of La4:La5 under both homeostatic and allogeneic stimulated conditions, we next investigated whether these drugs directly influenced FRC-derived laminins, given the crucial role of FRC in modulating laminin expression ^14,20^. We utilized two laminin KO mouse strains: FRC-Laminin4-KO mice (Pdgfrb-Cre+/− × La4^fl/fl^) have the laminin α4 gene specifically deleted in FRCs, and FRC-Laminin5-KO mice (Pdgfrb-Cre+/− × La5^fl/fl^) have the laminin α5 gene specifically deleted in FRCs ^14,52^. FRC-Lama4-KO and FRC-Lama5-KO mice were treated with MMF or FTY for 3 days, to specifically examine whether FRC-derived laminin α4 and α5 played a role in immunosuppressant-mediated effects on LN stromal organization.

In mLNs, as we observed in WT mice, 3-day treatment of MMF or FTY did not affect laminin ratios (**Figure 3F-I, Supplemental Figure 6C, D**). In FRC-Lama4-KO mice, MMF and FTY also did not affect laminin ratios (**Figure 6A-C, M**). However, in FRC-Lama5-KO mice, both drugs decreased the La4:La5 in CR of mLNs (**Figure 6D-F, M**). These results suggested that MMF and FTY effects on laminin composition were dependent on the FRC-derived laminin expression, indicating its critical role in maintaining the laminin balance in the mLN. The removal of FRC-derived laminin α5 appeared to unmask an early pro-inflammatory impact of MMF and FTY that was otherwise absent. This indicated that laminins specifically from FRCs acted as a protective factor in preserving stromal homeostasis in the mLNs, counteracting the early disruptive effects of these immunosuppressants.

**Figure 6.**
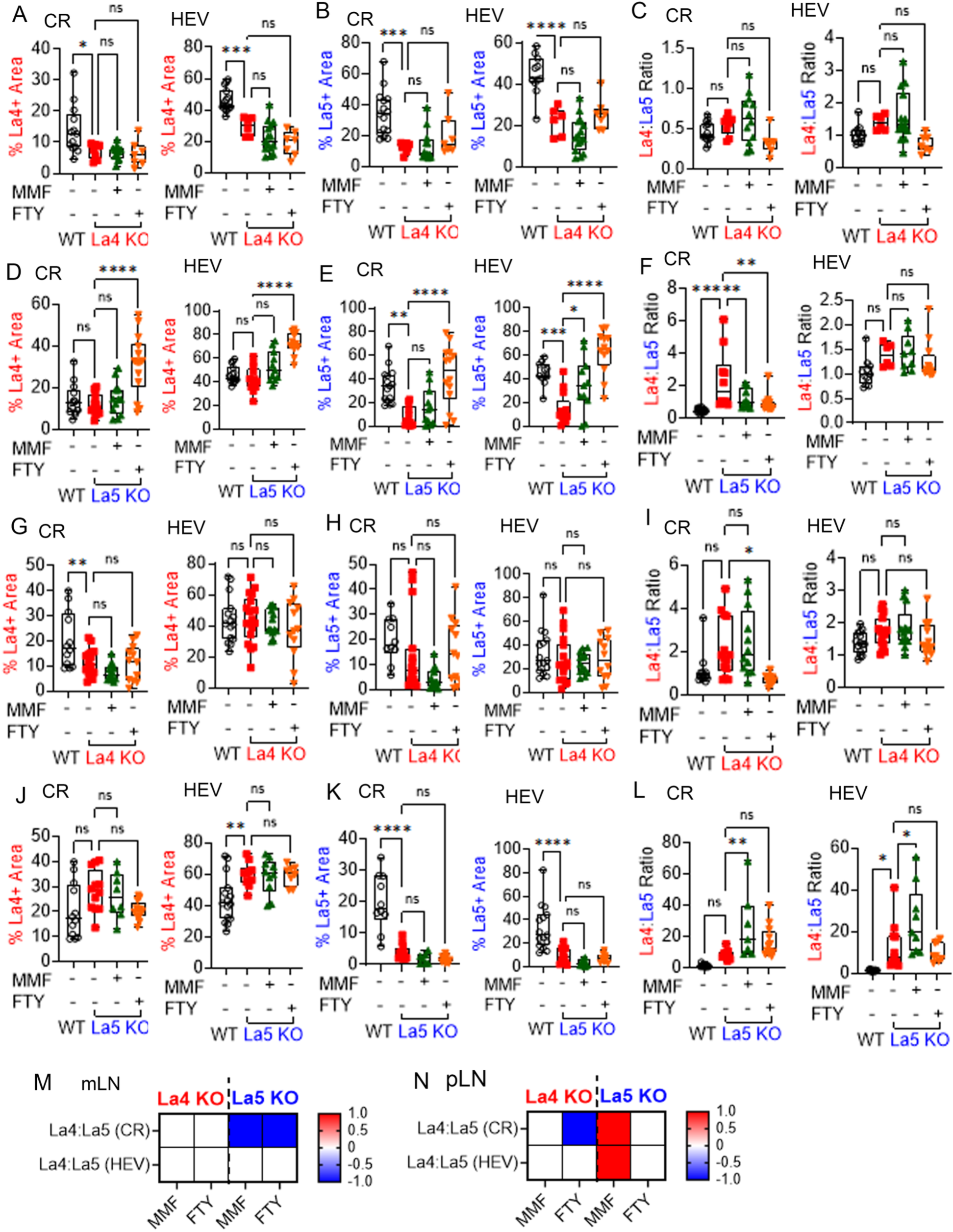
MMF and FTY decrease the La4:La5 in FRC-Lama5-KO mLNs and pLNs. IHC showing **A, D, G, J**) laminin α4; **B, E, H, K**) laminin α5; and **C, F, I, L**) La4:La5 in mLNs **(A-F, M)** and pLNs **(G-L, N)** of WT and FRC-Lama4-KO (**A-C and G-I**) and FRC-Lama5-KO mice (**D-F and J-L**) on day 3. **M, N**) Heatmap summarizes significant changes relative to controls in **M)** mLN and **N)** pLN. 3-5 mice/group. 2-3 LNs/mouse, 2-3 sections/LN, 7-30 fields/tissue. One-way ANOVA: * p < 0.05, ** p < 0.01, *** p < 0.001, **** p < 0.0001.

In pLNs, as we observed in WT mice, 3-day treatment of WT mice with MMF or FTY each increased La4:La5 (**Figure 4G-J, Supplemental Figure 7C, D**). MMF also increased La4:La5 in both CR and HEV of FRC-Lama5-KO mice (**Figure 6J-L, N**), but not in FRC-Lama4-KO mice (**Figure 6J-L, N**). In contrast, FTY decreased La4:La5 in CR of FRC-Lama4-KO mice (**Figure 6G-I, N**) but not in FRC-Lama5-KO mice (**Figure 6J-L, N**). These findings indicated that MMF and FTY modulated La4:La5 through distinct mechanisms that depended on the presence of laminins α4 and α5 from FRCs, although the precise pathways involved remain to be determined. The analysis of these KO models revealed that MMF relied on laminin α4 to increase laminin ratios, as its absence in FRC-Lama4-KO mice abolished the increase in La4:La5, while laminin α5 deficiency did not. Conversely, FTY depended on laminin α5 to maintain the La4:La5 ratio, as its absence revealed a decrease. These results demonstrated the distinct roles of laminins α4 and α5 in mediating immunosuppressant effects by remodeling LN architecture under MMF or FTY treatment. Collectively, these findings suggested a broader mechanistic link between immunosuppressive drugs and immune regulation, extending beyond their direct cellular effects to include the critical role of FRC-derived laminins in maintaining LN homeostasis

## Discussion

The gut plays as a critical role in maintaining transplant tolerance by serving as a key interface between the immune system and environmental stimuli ^53–55^. In this study, we performed an in-depth characterization of the previously underappreciated, complex interplay of immunosuppressants and several gut-associated processes, including the microbiome and metabolism, intestinal gene expression, mesenteric and peripheral LN organization, and lymphocyte trafficking over time. Though not designed to target the gut, our findings revealed that immunosuppressants exert profound, time-dependent effects on the gut environment. While early treatment exhibits distinct, drug-specific effects, prolonged immunosuppressant use leads to a convergent dysbiotic state in the gut microbiome by day 30. A particularly striking observation is the convergent enrichment of the *Muribaculaceae* family across all four drugs after long-term treatment. These bacteria harbor a repertoire of enzymes capable of efficiently breaking down complex carbohydrates and proteins ^42–44^, providing them with a competitive edge in an altered gut environment under immune suppression. Their metabolic versatility likely enables them to thrive under conditions of disrupted gut homeostasis, implying that sustained immunosuppression selectively favors organisms adapted to such changes. Furthermore, recent work identifying *Muribaculaceae* as pathobionts capable of inducing diabetes ^45^ offer a potential mechanistic link between prolonged immunosuppression and the metabolic complications often seen in transplant patients ^56–58^. This suggests that extended immune suppression may trigger a common mechanism affecting microbial communities.

Immune suppressants also triggered a common mechanism affecting epithelial and mucosal immunity (illustrated in **Figure 7**). All immunosuppressants disrupt antigen-driven B cell activation. Previous studies reported glucocorticoids directly suppress B cell function by inhibiting NF-κB-dependent transcription, leading to reduced expression of cytokines, costimulatory molecules, and survival factors essential for B cell activation and differentiation ^59,60^. Calcineurin inhibitors impair NFAT-dependent signaling by blocking dephosphorylation of NFAT proteins, thereby inhibiting transcriptional programs involved in B cell proliferation and antibody production while also inhibiting helper T cells ^61^. In contrast, mTOR inhibitors and anti-metabolites interfere more indirectly by suppressing T follicular helper (Tfh) cell development and function, leading to impaired T cell-dependent B cell help, germinal center formation, and class-switch recombination ^62^. While mucosal adaptive immunity is predominately suppressed, the upregulation of a class-switched mechanism, as in Ighg2b gene activation, suggests selective survival or expansion of mucosal-associated or microbiota-reactive B cell subsets. This is supported by epithelial gene expression patterns, showing early induction of circadian resetting program and xenobiotic detoxification pathways, followed by late activation of antimicrobial peptides, ROS-generating oxidases, and stress-responsive enzymes. These changes point to a compensatory host mechanism to counteract potential pathobiont expansion and epithelial stress. However, prolonged treatments still led to transcriptomic changes in epithelial regenerative capacity, lipid metabolism, and structural remodeling. These impaired intestinal functions may predispose the mucosa to vulnerability during sustained immunosuppressive therapy. Overall, these findings reveal a complex interplay between immunosuppressive drug disturbance and the host’s attempt to preserve mucosal homeostasis. Further experiments to pinpoint the precise intestinal responders common to all immune suppressants classes will be essential to understand off-target drug effects and to guide strategies to mitigate complications.

**Figure 7.**
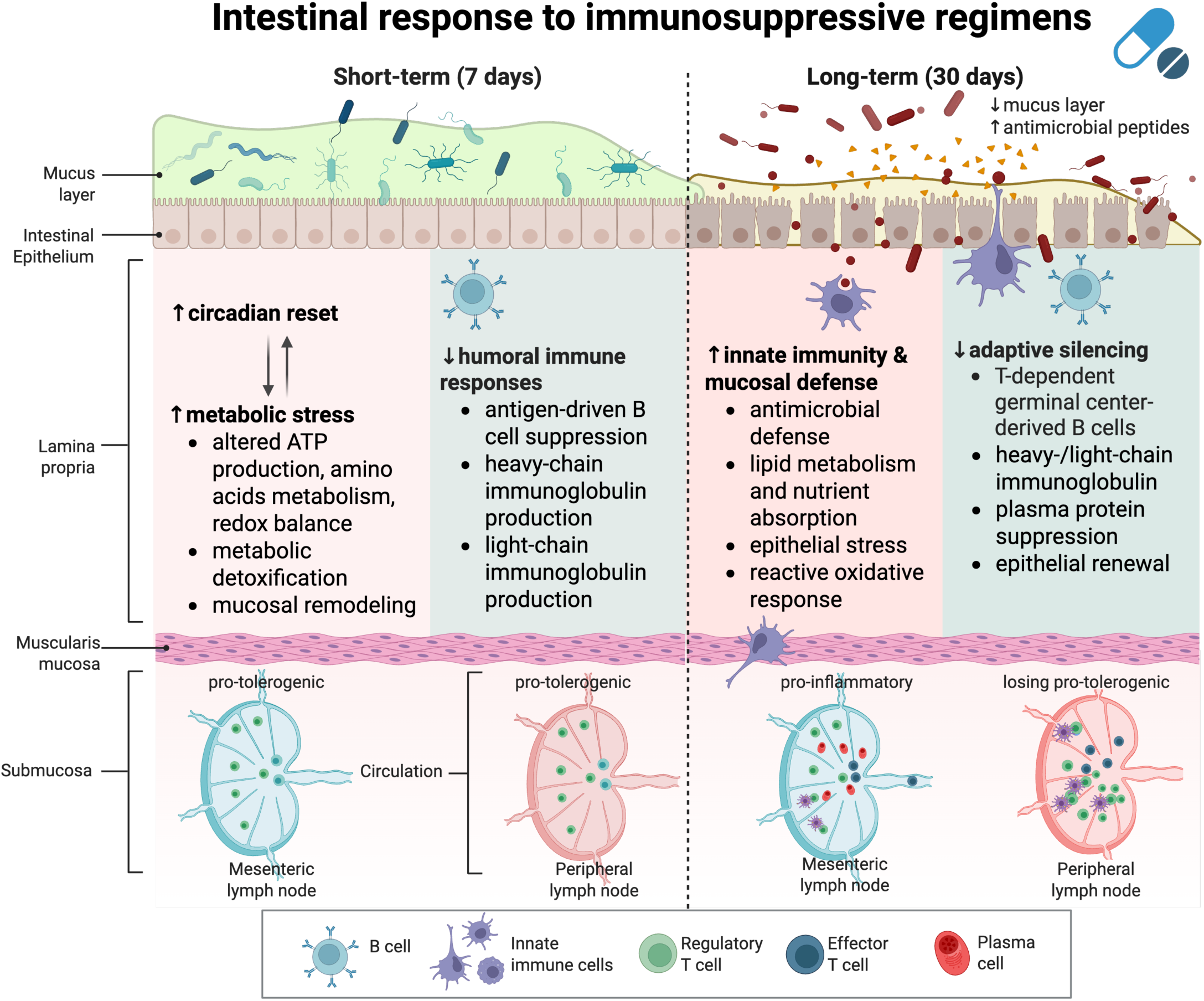
Illustration of shared intestinal responses in response to immunosuppressant regimens. Created in BioRender. Ma, B. (2025) https://BioRender.com/ctcfgx9.

Beyond shared microbial and intestinal shifts, each immunosuppressant exhibited distinct intestinal and metabolic signatures that revealed both intended and unintended mechanistic insights. By day 30, intestinal gene expression profiles became increasingly broad, underscoring the tissue’s heightened susceptibility to off-target drug effects. These differences likely arise from a combination of direct pharmacologic actions on host epithelial and immune cells, compensatory host responses, and complex interactions with the gut microbiota. At the metabolic level, each drug elicited shared and unique luminal metabolomic changes. TAC broadly altered amino acid metabolism and reduced levels of microbial-derived metabolites such as SCFAs, consistent with its known gastrointestinal side effects such as diarrhea and nonspecific abdominal discomfort ^63^. PRED exerted a widespread suppressive effect on metabolism, consistent with its ability to inhibit cellular functions and pro-inflammatory cytokines ^64^. This resulting reduction in amino acid derivatives, SCFAs, TMA, and polyamines provides insights into its side effects, including diabetes and osteoporosis ^47–49^. MMF targeted purine metabolism via IMPDH inhibition and suppressed the TCA cycle and branched-chain amino acid (BCAA) pathways, impairing ATP production and enterocyte protein synthesis. This may underlie its distinct profile of gastrointestinal toxicity and inflammation ^65,66^, compromised barrier integrity ^36^, and increased epithelial apoptosis and villous atrophy ^67^. FTY, in contrast, produced the mildest metabolic disruption, primarily affecting nucleotide turnover and polyamine biosynthesis, a profile that may explain its generally better tolerability ^68^.

Despite their distinct profiles, all immunosuppressants consistently affected amino acid metabolism and energy processing, suggesting a fundamental mechanism underlying long-term immunosuppression. The reduction in SCFA-producing bacteria can disrupt metabolic processes such as blood pressure regulation, lipogenesis, and blood glucose control ^69^. Studies have demonstrated that probiotics can reverse conditions like tacrolimus-induced gut dysbiosis, MMF-related intestinal toxicity and inflammation, SCFA depletion, and hypertension by restoring the vascular redox state and improving endothelial nitric oxide synthase (eNOS) function ^69–72^. On the other hand, differences in how these drugs alter amino acid metabolism indicate that each exerts distinct influences on protein synthesis and cellular energy balance, which may explain their varying effects on different immune cell populations. Arginine metabolism, for example, is critical because it affects eNOS activity through the arginine-NO pathway, influencing NO production and ROS generation, changes that can impair endothelial function and modulate immune cell activation ^73^. Both calcineurin and mTOR inhibitors drastically affect amino acid regulation ^31^, including arginine metabolism, which aligns with their documented vascular complications ^74–76^ and supports targeted probiotic therapies to mitigate adverse metabolic and vascular outcomes. Ultimately, the shared and drug-specific intestinal and metabolic signatures represent actionable targets for monitoring and mitigating off-target toxicity, offering new opportunities to improve safety and precision in transplantation medicine.

The impact of immunosuppressants on lymphoid tissue organization further illustrates the complex nature of their effects. Our data revealed anatomically distinct and temporally dynamic responses with lymphoid tissues. pLNs maintained an anti-inflammatory state, evidenced by elevated Tregs and increased La4:La5 ratios, particularly during early treatment phases. This response is consistent with the intended immunosuppressive action of these drugs. In contrast, mLNs, which drain the intestine, eventually developed a pro-inflammatory environment, characterized by reduced Tregs and decreased La4:La5. This compartment-specific regulatory effect may underlie some of the gastrointestinal side effects observed with immunosuppression ^32^. As illustrated in **Figure 7**, these changes likely reflect feedback from the intestinal microenvironment, where metabolic stress and mucosal adaptation shape lymphoid architecture. Further insights into LN FRC-derived laminins α4 and α5 in mediating these effects suggest that stromal responses play a key role in mediating these effects ^14^. Notably, MMF and FTY both preserved tolerogenic LN architecture during allostimulation through the dual regulation of Tregs and stromal organization, though they were distinctly dependent on laminin α4 or laminin α5. These findings highlight a critical gap in our understanding of how immunosuppressive drugs precisely affect the gene expression, function, and phenotype of LNSCs, and they emphasize the need for further research to develop more precise therapeutic strategies.

This study does have several limitations. Our 30-day mouse studies may not fully capture the complexity of long-term immunosuppression in human transplant patients, and extended longitudinal studies are needed to better mirror clinical outcomes. Moreover, we evaluated single drugs rather than the combination regimens commonly used in clinical practice. Additionally, although our allogeneic stimulation model provided valuable insights into transplant-related alloimmune responses, the absence of an allograft precluded the pro-inflammatory context typical of solid organ transplantation, potentially underestimating the drug impact on immune regulation during active alloimmune responses. Nonetheless, our study established fundamental insights into immunosuppressants effects, and revealed an underappreciated while profound impacts on the intestine microenvironment, providing a critical basis for future transplant investigations. Overall, our findings highlight an understudied aspect of immunosuppressant effects on the intestine and lymphoid tissue, emphasizing the need for strategies such as conserved biomarker-monitoring, microbiome-targeted interventions, and tissue-specific drug delivery to enhance therapeutic precision and reduce side effects.

## Methods and Materials

### Study approval

All procedures, as described previously ^31,32,77,78^ involving mice were performed in accordance with the guidelines and regulations set by the Office of Animal Welfare Assurance of the University of Maryland School of Medicine under the approved IACUC protocol nos. 05150001, 0318002, 1220001, and AUP-00000397.

### Reagents

USP grade immunosuppressants were used, when available. All drugs were prepared in a manner approved by the University of Maryland School of Medicine IACUC: TAC (USP grade, MilliporeSigma, Burlington, MA) was reconstituted in DMSO (USP grade, MilliporeSigma) at 20 mg/ml for storage. At time of use, TAC/DMSO stock was diluted with absolute ethanol (USP grade, Decon Labs, King of Prussia, PA) to 1.5 mg/ml then further diluted 1:5 in sterile PBS and injected at 10 μl/gm s.c. (3 mg/kg/day) ^30^. PRED (MilliporeSigma) was reconstituted in PBS at 50 mg/ml for storage and then diluted to 0.5 mg/ml in PBS and injected at 5mg/kg/d i.p. ^79^. MMF (MilliporeSigma) was reconstituted at 30 mg/ml in DMSO for storage and then diluted in PBS to 1.5 mg/ml and at 30 mg/kg/d i.p. ^80^. FTY (Fingolimod, MilliporeSigma) was reconstituted at 3 mg/ml in 1:1 PBS:Ethanol for storage and then diluted 1:10 in PBS and administered at 3 mg/kg/d p.o.^81^. The control group received PBS only.

### Mice experiments

Mouse experiments were performed according to ARRIVE guidelines (https://arriveguidelines.org). 8- to 14-week-old female C57BL/6 mice were purchased from The Jackson Laboratory (Bar Harbor, ME, USA). Mice were maintained at the University of Maryland School of Medicine Veterinary Resources animal facility. Only female mice were used to ensure a high degree of homogeneity within our study groups. The Pdgfrb-Cre+/– x La4fl/fl ^52^ and Pdgfrb-Cre+/– x La5fl/fl ^14^ conditional knockout (KO) mice were previously developed in our laboratory. All mice were cohoused for a minimum of 2 weeks prior to experiments to normalize microbiota. During this period, mice from different treatment groups were housed in the same room and adjacent cages under identical conditions to ensure normalized environmental exposure. At experimental day 0, the mice were randomly separated to different experimental groups, and then kept groups in separate cages to prevent cross exposure. Mice received daily immunosuppression following the dosages in **Table 2**. In allogeneic stimulation experiment, mice received 10^7 fully allogeneic BALB/c splenocytes intravenously (i.v.) on day 0 followed by drug treatment. On the day of harvest, the mice were euthanized by CO_2_ narcosis, intraluminal stool samples collected for metabolomic and microbiome analyses, cardiac puncture utilized for blood collection, and mesenteric and peripheral (axillary, inguinal, and brachial) LNs and portions of the small intestine between the duodenum and jejunum harvested for immunological assays. All procedures involving mice were performed in accordance with the guidelines and regulations set by the Office of Animal Welfare Assurance of the University of Maryland School of Medicine.

**Table 2.** Immunosuppression drugs and dosage.

| Treatment Groups | Dose*(mg/kg/day) | Source (catalog #) | Route |
| --- | --- | --- | --- |
| **Tacrolimus** | 3 mg/kg/day | Sigma Aldrich (Y0001926) | Subcutaneous <sup>100</sup> |
| **Prednisolone** | 5 mg/kg/day | Sigma Aldrich (P6004) | Intraperitoneal <sup>79</sup> |
| **Mycophenolate mofetil** | 30 mg/kg/day | Sigma Aldrich (SML0284) | Intraperitoneal <sup>101</sup> |
| **FTY** | 3 mg/kg/day | Sigma Aldrich (SML0700) | Oral <sup>102</sup> |

### Stool specimen collection, DNA extraction, and metagenomic sequencing

As described previously ^31,32,77,78^, stool samples were collected and stored immediately in DNA/RNA Shields (Zymo Research, Irvine, CA, USA) at −80°C. This stabilizes and protects the integrity of nucleic acids and minimizes the need for immediate processing of specimens. The Quick-DNA Fecal/Soil Microbe kit (Zymo Research, Irvine, CA, USA) was used to extract DNA from 0.15-0.25 grams of fecal samples. To ensure that no exogenous DNA contaminated the samples, we included negative extraction controls. Construction of the metagenomic sequencing libraries was performed as previously ^31,32,78^. In brief, the Nextera XT Flex Kit (Illumina), according to the manufacturer’s recommendations, libraries were then pooled together in equimolar proportions before they were sequenced on a single Illumina NovaSeq 6000 S2 flow cell at Maryland Genomics at the University of Maryland School of Medicine. 92.7±8.5 (mean±s.e.) million reads per sample were obtained.

### Gut microbiome analyses

Sequence analyses were performed as previously described ^31,32,77,78^. Metagenomic sequence reads mapping to Genome Reference Consortium Mouse Build 39 of strain C57BL/6J (GRCm39) were removed using BMTagger v3.101 ^82^. Sequence read pairs were removed when one or both the read pairs matched the genome reference. The Illumina adapter was trimmed and quality assessment was performed using default parameters in fastp (v.0.21.0) ^83^. The taxonomic composition of the microbiomes was established using Kraken2 (v.2020.12) and Braken (v. 2.5.0) using the comprehensive mouse microbiota genome catalog ^84^. Phyloseq R package (v1.38.0) was used to generate the barplot and diversity index. Linear discriminant analysis (LDA) effect size (LEfSe) analysis was used to identify fecal phylotypes that could explain the differences. The α value for the non-parametric factorial Kruskal-Wallis (KW) sum-rank test was set at 0.05 and the threshold for the logarithmic LDA model score for discriminative features was set at 2.0. An all-against-all BLAST search was performed in the multiclass analysis. Taxonomic ordination graphs were created with the microViz (v0.12.4) ^85^. The metagenomic dataset was mapped to the protein database UniRef90 to ensure the comprehensive coverage in functional annotation, and was then summarized using HUMAnN3 (Human Microbiome Project Unified Metabolic Analysis Network) (v0.11.2) to determine the presence, absence, and abundance of metabolic pathways in a microbial community. MetaCyc pathway definitions and MinPath were used to identify a parsimonious set of pathways summarized in HUMAnN3 in the microbial community. Canonical Correspondence Analysis (CCA) was performed using the vegan package ^86,87^ based on the Bray-Curtis distance. Based on their eigenvalues, CA1 and CA2 were selected as the major components. To evaluate microbiome profile differences across groups and time points, the line plot was generated using the average pairwise Bray-Curtis distances between all group pairs at each time point.

### Metabolite extraction and metabolome analyses

As previously described ^31,32,77,78^, capillary electrophoresis-mass spectrometry (CE/MS) was used for measuring metabolome of intraluminal stool (luminal/local) to obtain a comprehensive quantitative survey of metabolites (Human Metabolome Technologies, Boston, MA, USA) for immunosuppressants TAC, PRED, MMF, and a no-treatment control group processed concurrently. For FTY, stool pellets were collected at the same time points for metabolomic analysis due to technical limitations in obtaining sufficient sample volumes for comprehensive profiling. The no-treatment control for FTY also used stool pellets and processed concurrently. All subsequent analyses were performed using the respective no-treatment control groups and the same specimen types, processed concurrently with their corresponding experimental groups, to ensure valid comparisons. ∼10-30 mg of stool was weighed at the time of collection using a company-provided vial and stored at −80°C until shipped to vendor on dry ice. QC procedures included standards, sample blanks and internal controls that were evenly spaced among the samples analyzed. Compound identification was performed using a CE/MS library of >1,600 annotated molecules.

To reduce the influence of measurement noise, rigorous data pretreatment was performed according to validated procedures ^88^ and are as previously described ^31,32,78^. Metabolites were annotated using PubChem ^89^, KEGG ^90^, and HMDB ^91^ annotation frameworks that leverage cataloged chemical compounds, known metabolic characterization, and functional hierarchy (i.e., reaction, modules, pathways). Partial least squares discriminant analysis (PLS-DA) implemented using mixOmics (vers. 6.18.1) was used ^92^. The “sparseness” of the model was adjusted by the number of components in the model and the number of variables within each component based on the classification error rate with respect to the number of selected variables. Tuning was performed one component at a time, and the optimal number of variables to select was calculated. The volcano plot combines results from fold change (FC) analysis to show significantly increased metabolites after 7-day tacrolimus treatment. A metabolite is shown if FC is >2 and the p-value is <0.05 based on 2-sample t-tests. Original metabolite measurements without normalization were used in the FC analysis. For the metabolites annotated in a specific functional pathway, metabolite set enrichment analysis (MSEA) was performed ^93^.

### RNA isolation, transcriptome sequencing, and bioinformatics analyses

RNA isolation, transcriptome sequencing, and bioinformatics analyses are as previously described ^31,32,78^. Dissected intestinal tissues were stored immediately in RNAlater solution (QIAGEN) in RNAse-free tubes (Denville Scientific, Holliston, MA) to stabilize and protect the integrity of RNA ^94^. Specimens were stored at −80°C until extraction. For each sample, total RNA was extracted from ∼1 cm of ileum. Prior to the extraction, 500 µl of ice-cold RNase free PBS was added to the sample. To remove the RNAlater, the mixture was centrifuged at 8,000x*g* for 10 min and the resulting pellet resuspended in 500 µl ice-cold RNase-free PBS with 10 µl of β-mercaptoethanol. A tissue suspension was obtained by bead beating procedure using the FastPrep lysing matrix B protocol (MP Biomedicals, Solon, OH) to homogenized tissues. RNA was extracted from the resulting suspension using TRIzol Reagent (Invitrogen, Carlsbad, CA) following the manufacturer’s recommendations and followed by protein cleanup using Phasemaker tubes (Invitrogen) and precipitation of total nucleic acids using isopropanol. RNA was resuspended in DEPC-treated DNAase/RNAase-free water. Residual DNA was purged from total RNA extract by treating once with TURBO DNase (Ambion, Austin, TX, Cat. No. AM1907) according to the manufacturer’s protocol. DNA removal was confirmed via PCR assay using 16S rRNA primer 27 F (5′-AGAGTTTGATCCTGGCTCAG-3′) and 534 R (5′-CATTACCGCGGCTGCTGG-3′). The quality of extracted RNA was verified using the Agilent 2100 Expert Bioanalyzer using the RNA 1000 Nano kit (Agilent Technologies, Santa Clara, CA). Ribosomal RNA depletion and library construction were performed using the RiboZero Plus kit and TruSeq stranded mRNA library preparation kit (Illumina) according to the manufacturer’s recommendations. Libraries were then pooled together in equimolar proportions and sequenced on a single Illumina NovaSeq 6000 S2 flow cell at the Genomic Resource Center (Institute for Genome Sciences, University of Maryland School of Medicine) using the 150 bp paired-end protocol. 143,2±53.4 (mean±s.e.) million reads per sample were obtained. The quality of sequencing was evaluated by using FastQC ^95^. Reads were aligned to the mouse genome (Mus_musculus.GRCm39) using HiSat (version HISAT2-2.0.4) ^96^ and the number of reads that aligned to the coding regions was determined using HTSeq ^97^. Significant differential expression was assessed using DESeq2 ^98^ with an FDR value ≤ 0.05 and FC>2. The over-representative analysis was done by importing Differentially Expressed Genes (DEGs) against GO ontologies using the enrichGO function of clusterProfile Bioconductor package ^99^. Only the ontology terms with q value <0.05 were used for plotting. The normalized enrichment score was calculated as the ratio of the proportion of DEGs annotated to a given GO term to the proportion of genes in the reference annotated to the same GO term. UpSet plots were generated using UpSetR (vers. 1.4.0) to analyze overlapping DEGs between days 7 and 30 and across different treatments.

### Flow cytometry

The procedure is as previously described ^31,32^. To produce single-cell suspensions, LNs were disaggregated and passed through 70-μm nylon mesh screens (Thermo Fisher Scientific, Waltham, MA). Antibodies against surface molecules were incubated with cell suspensions for 30 min at 4°C (**Supplemental Table 4**) then washed with FACS buffer [PBS with 0.5% w/v bovine serum albumin] twice. Cells were then permeabilized, according to manufacturer’s protocol, with Foxp3/Transcription Factor Staining Buffer Set (eBioscience, San Diego, CA), washed with FACS buffer, and subsequently stained with antibodies for intracellular molecules at 4°C. Samples were analyzed with an LSR Fortessa Cell Analyzer (BD Biosciences), and data were analyzed using FlowJo software version 10.6 (BD Biosciences). Single color controls (cells stained with single surface marker antibody) and unstained controls were used for flow channel compensation.

### Immunohistochemistry

We excised mLN, pLN and segments of the intestine between the duodenum and jejunum then immediately submerged in OCT compound (Scigen Paramount, CA, USA). Cryosections (6-7 μm for LNs, 10 μm for intestine) were cut in duplicate or triplicate depending on size of block using a Microm HM 550 cryostat (ThermoFisher Scientific). Slides were then fixed in cold 1:1 acetone:methanol for 5 min, air dried for 15 min, and stored at −80°C until use. At the time of staining, slides were thawed for 15 minutes at room temperature then sections were rehydrated in PBS and blocked with 10% secondary antibody host serum PBST (PBS + 0.03% Triton-X-100 + 0.5% BSA). The slides were then stained for 1 hour with primary antibodies at room temperature (diluted 1:50-1:200 in PBST) for 1 hour incubated with secondary antibodies (diluted 1:100-1:400 in PBST) for 1 hour. After staining, slides were fixed with 4% paraformaldehyde in PBS for 5 min, quenched with 1% glycerol in PBS for 5 min, and mounted with Prolong Gold Antifade Mountant with or without DAPI (Thermo Fisher Scientific). Images were acquired using Accu-Scope EXC-500/Nikon (Accu-Scope, Commack, NY, and Nikon, Tokyo, Japan) and analyzed using Volocity software (Quorum Technologies Inc., Sacramento, CA). The antibodies used are listed in **Supplemental Table 4**. For each mouse, 1-2 mLNs and 2-3 pLNs, and 1 piece of intestine were collected. All samples from each treatment group were combined and analyzed as a block, 2-3 sections/block were placed on slides, and 7-30 fields/slide were analyzed. The average of MFI for each treatment group was calculated by averaging the MFI values across all slides from all mice, within demarcated HEVs and CR regions of mLN and pLN and whole intestinal images. Qualitative heat maps were generated (GraphPad prism) to express changes in IHC marker expression level relative to control using −1 to represent “decreased” (blue in the heatmap), 0 to represent “unchanged” (white in the heatmap), and 1 to represent “increased” (red in the heatmap). GraphPad Prism 10.3.1 (San Diego, CA, USA) was used for analysis with statistical significance defined as *P* < 0.05. Tukey’s multiple comparison tests of one-way ANOVA were used for comparisons of fluorescence.

## Declarations

### Ethics Approval and Consent to Participate

All experimental procedures involving mice were approved by and performed in accordance with the guidelines and regulations of the Institutional Animal Care and Use Committee (IACUC) of the University of Maryland School of Medicine under the approved protocol numbers 05150001, 0318002, 1220001, and AUP-0000397.

### Consent for Publication

Not applicable.

### Availability of data and materials

The authors confirm that the data supporting the findings of this study are available within the article and indicated supplementary materials. The data that support the findings of this study are openly available; metagenome sequences were submitted to GenBank under BioProject PRJNA809764 (https://www.ncbi.nlm.nih.gov/bioproject/PRJNA809764) with the SRA study ID SRP361281. RNA-Seq data were deposited in the NCBI Gene Expression Omnibus database (GSE288074). The preprint of this study is available at https://doi.org/10.1101/2025.01.02.631100.

### Competing interests

No potential financial and non-financial conflict of interest was reported by the authors.

### Funding Declaration

This work was supported by the National Institute of Health (NIH) National Heart, Lung and Blood Institute award R01HL148672 (JSB/BM), NIH National Institute of Allergy and Infectious Diseases U01AI170050 (BM/JSB/VM), R01AI114496 (JSB) and training grant T32AI95190-10 (SJG). University of Maryland Baltimore ORD-OTT No. BM-2025-077.

### Authors’ contributions

B. M., L. W., and J. S. B. designed the experiments. L. W., A. K., S. J. G., D. K., L. L., R. L., V. S., W. P., M. W., and B. L. conducted and analyzed the mice experiments. H. W. L. performed nucleic acid extraction and sequencing library preparation. B. M. and Y. S. performed the bioinformatics analyses. B. M., L. W., A. K., S. J. G., M. F., Y. S., and J. S. B. wrote the manuscript. B. M. and J. S. B. are co-corresponding authors for this manuscript.

## Notes

### Competing Interest Statement

The authors have declared no competing interest.

### Summary of Updates

We analyzed DEGs common to all immunosuppressant treatments to identify shared drug-induced effects, which was not in previous version. The results are significant part of the paper. Clustering analysis of these conserved DEGs at days 7 and 30 revealed distinct temporal expression patterns, particularly affecting genes involved in mucosal immunity, epithelial homeostasis, metabolism, and circadian regulation.

