## Supplementals for "Immunosuppressants Rewire the Gut Microbiome-Alloimmune Axis Through Time-Dependent and Tissue-Specific Mechanisms"

#### Supplemental Figures

**Supplemental Figure 1: Canonical Correspondence Analysis (CCA) to demonstrate *de novo* clustering of gut microbiome** in **A**) taxonomic groups and **B**) microbial functional pathways based on Bray-Curtis distance. Taxonomic composition of the microbiomes using Kraken2 (v.2020.12) [1] and Braken (v. 2.5.0) [2] and the comprehensive mouse gut metagenome catalog (CMGM) [3]. Functional pathways characterized using HUMAnN3 (Human Microbiome Project Unified Metabolic Analysis Network) (v3.0.0.alpha.3) [4] and Uniref90 database[5].

**Supplemental Figure 2: Immunosuppressants induce distinct alterations in DC and Treg populations in the small intestine. A)** CD11c+ DCs and **B)** Tregs after 3, 7, and 30 days of drug treatment. **C)** Heatmap summarizes significant changes in DCs and Tregs compared to controls without drug treatment. Red increase, white no change, blue decrease. 3 mice/group, 3 pieces of intestine, 2-3 sections/intestine, 7-30 fields/tissue. One-way ANOVA: * $p < 0.05$, ** $p < 0.01$, *** $p < 0.001$, **** $p < 0.0001$.

**Supplemental Figure 3: Analysis of differentially expressed genes (DEGs)** between days 7 and 30 for each individual immune suppressants of **A**) TAC, **B**) PRED, **C**) MMF, and **D**) FTY720 using UpSet plots. Detailed list of genes is listed in **Supplemental Table 2B-E**. Horizontal bars (left) indicate total DEG numbers, vertical bars show the size of unique and overlapping gene sets, and dots below indicate the specific conditions for each intersection, and connected dots mean those sets are shared. Arrows indicate up- or down-regulation of DEGs.

**Supplemental Figure 4: Analysis of differentially expressed genes (DEGs) within and across immunosuppressant treatments.** UpSet plots showing intersections of DEGs for each treatment compared to control, including **A**) upregulated and **B**) downregulated DEGs. Detailed list of genes are listed in **Supplemental Table 2B-E**. Horizontal bars (left) indicate total DEG numbers, vertical bars show the size of unique and overlapping gene sets, dots below indicate the specific conditions for each intersection, and connected dots mean those sets are shared. Arrows indicate up- or down-regulation of DEGs. **C**) Functional enrichment analysis of intestinal transcriptional responses to immunosuppressant treatments. Dot plot illustrating over-represented immune-related functional pathways of DEGs in small intestinal tissues across different treatments compared to the control group, analyzed at 7 and 30 days of treatment. Gene Ontology pathways used. Dot size represents the -log10(p-value), indicating the statistical significance of pathway enrichment.

**Supplemental Figure 5: Gut luminal metabolites after immune suppressant drug treatment. A)** PLS-DA clusters by treatment groups. Circles indicate 95% confidence region. **B**) Metabolites contributing to separation of each treatment group in PLS-DA, ranked by variable importance in projection (VIP). **C**) Hierarchical clustering heatmap of metabolites after immune suppressant drug treatment. Specimens collected after treatment for 7 days, compared to no drug control. Top 50 features shown. Color bar indicates the scaled z-score of each feature. Ward linkage clustering based on Euclidian distance.

**Supplemental Figure 6: Immunosuppressants distinctively influence gut metabolism.** Volcano plots combine results from fold change (FC) analysis to show significantly increased metabolites after 7 days for **A**) TAC, **B**) PRED, **C**) MMF, or **D**) FTY treatment compared to no treatment control. A metabolite is shown if FC is >2 and p value is <0.05 based on 2-sample t-tests. Original metabolite measurement without normalization in FC analysis.

**Supplemental Figure 7: Effects of TAC, PRED, MMF, and FTY720 on mLN cell content, cell distribution, and structure.** IHC of **A)** CR and **B)** HEV Foxp3+ Tregs on days 3, 7, and 30 of drug treatment. IHC of **C)** CR and **D)** HEV La4:La5 on days 3, 7, and 30 of drug treatment. 3 mice/group 2-3 LNs/mouse, 2-3 sections/LN, 7-30 fields/tissue One-way ANOVA. * $p < 0.05$; ** $p < 0.01$, *** $p < 0.001$, **** $p < 0.0001$.

**Supplemental Figure 8: Effects of TAC, PRED, MMF, and FTY720 on pLN cell content, cell distribution, and structure.** IHC of **A)** CR and **B)** HEV Foxp3+ Tregs on days 3, 7, and 30 of drug treatment. IHC of **C)** CR and **D)** HEV La4:La5 on days 3, 7, and 30 of drug treatment. 3 mice/group, 2-3 LNs/mouse, 2-3 sections/LN, 7-30 fields/tissue. One-way ANOVA. * $p < 0.05$; ** $p < 0.01$, *** $p < 0.001$, **** $p < 0.0001$.

**Supplemental Figure 1.**

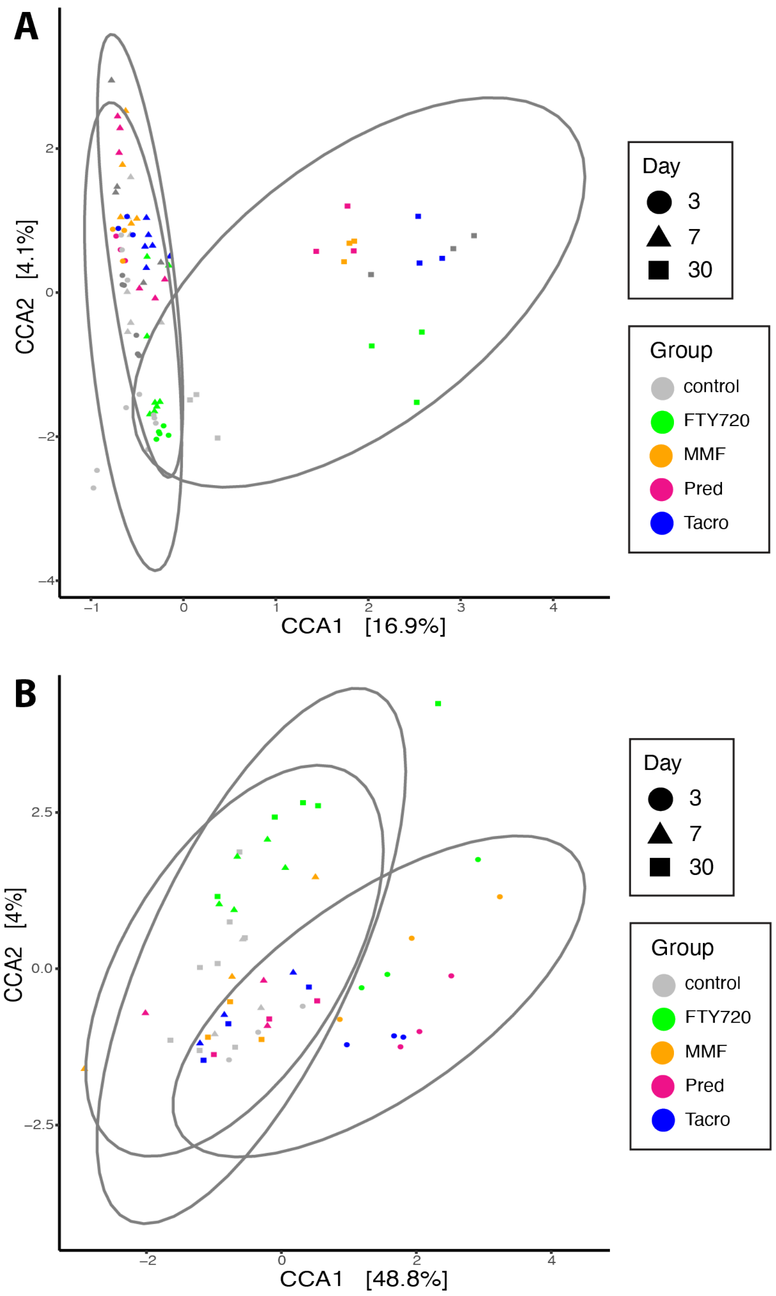

**Supplemental Figure 2.**

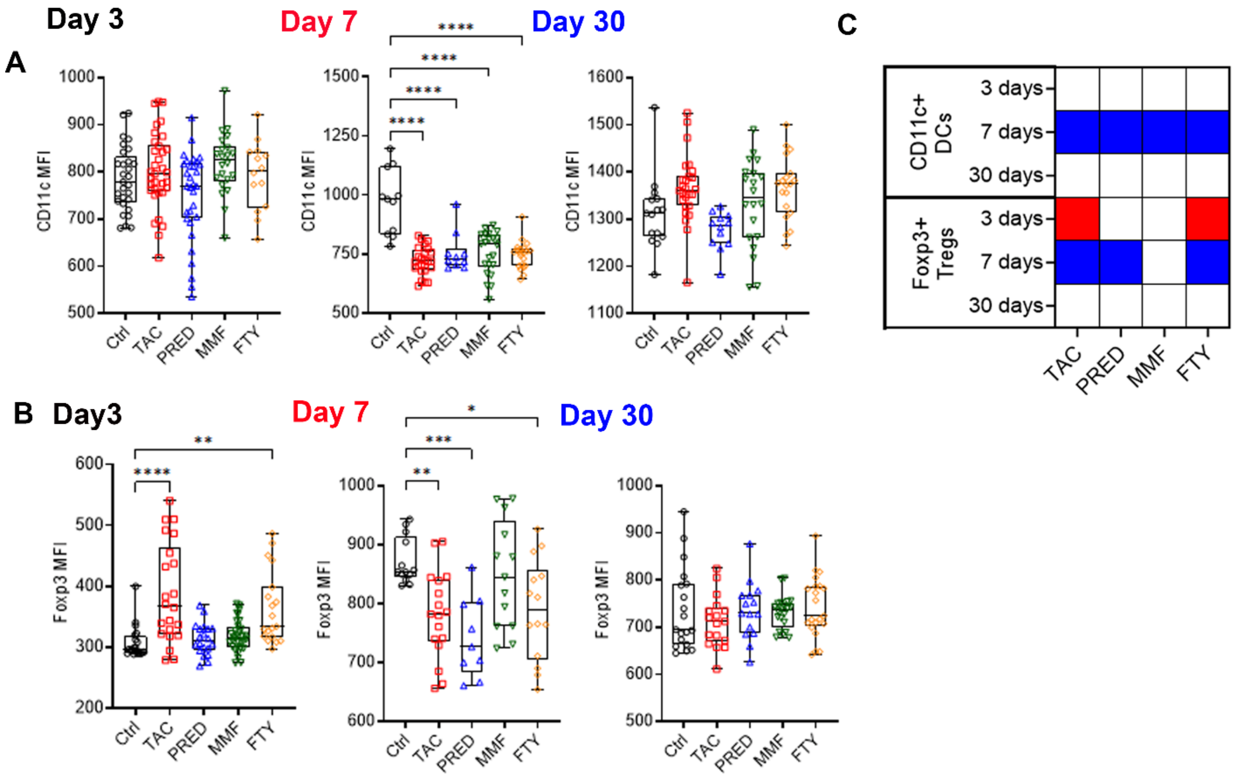

**Supplemental Figure 3. A)**

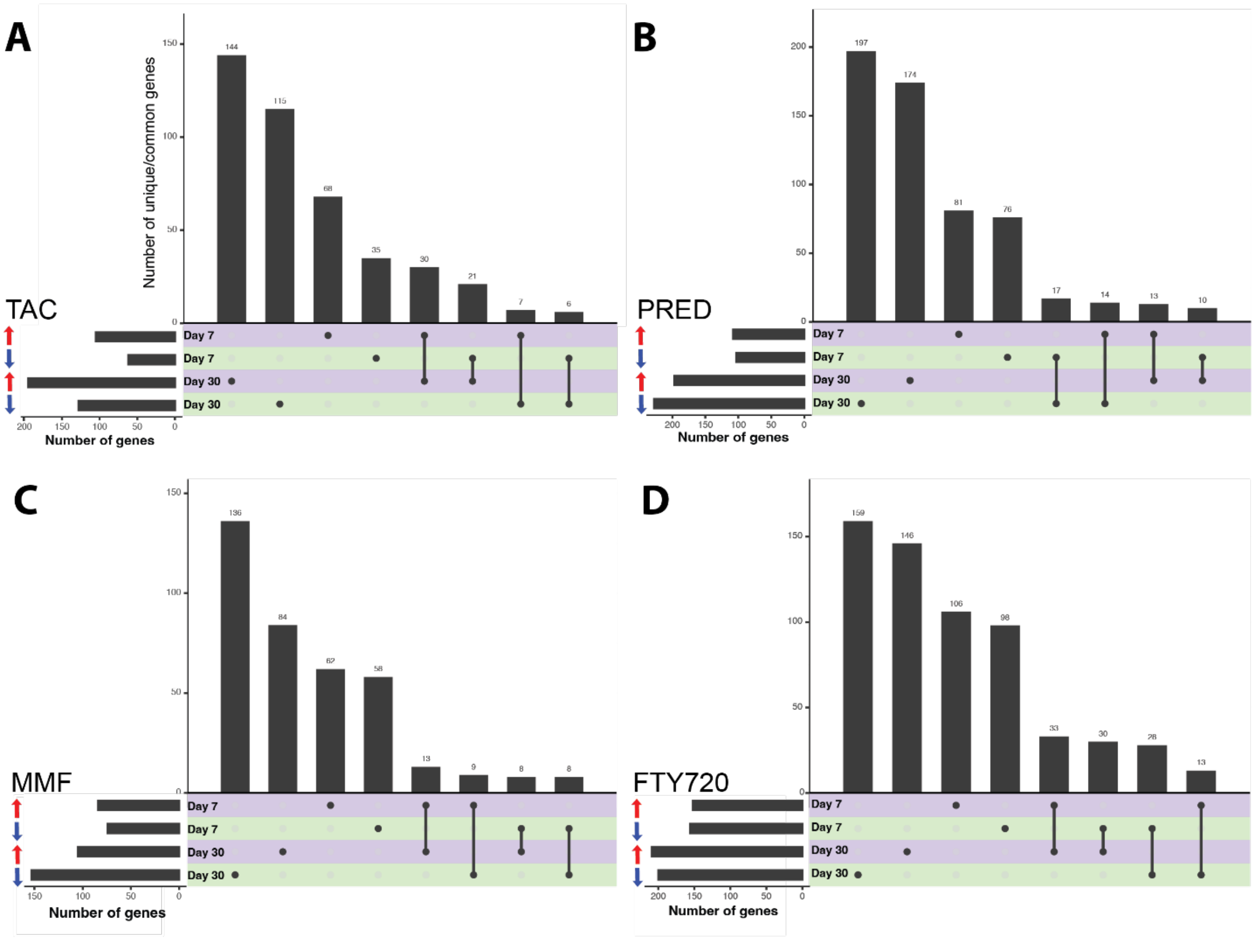

**Supplemental Figure 3.**

**A)**

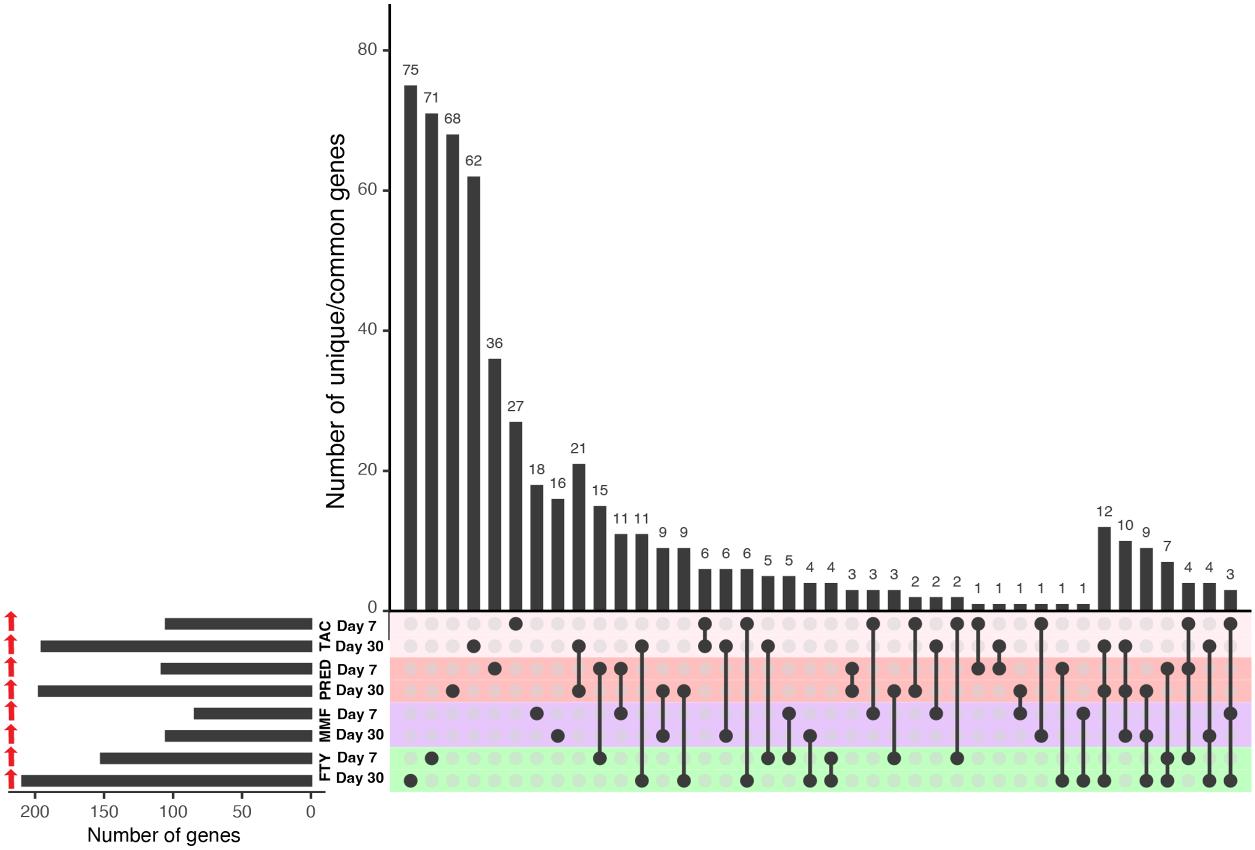

**Supplemental Figure 3.**

**B)**

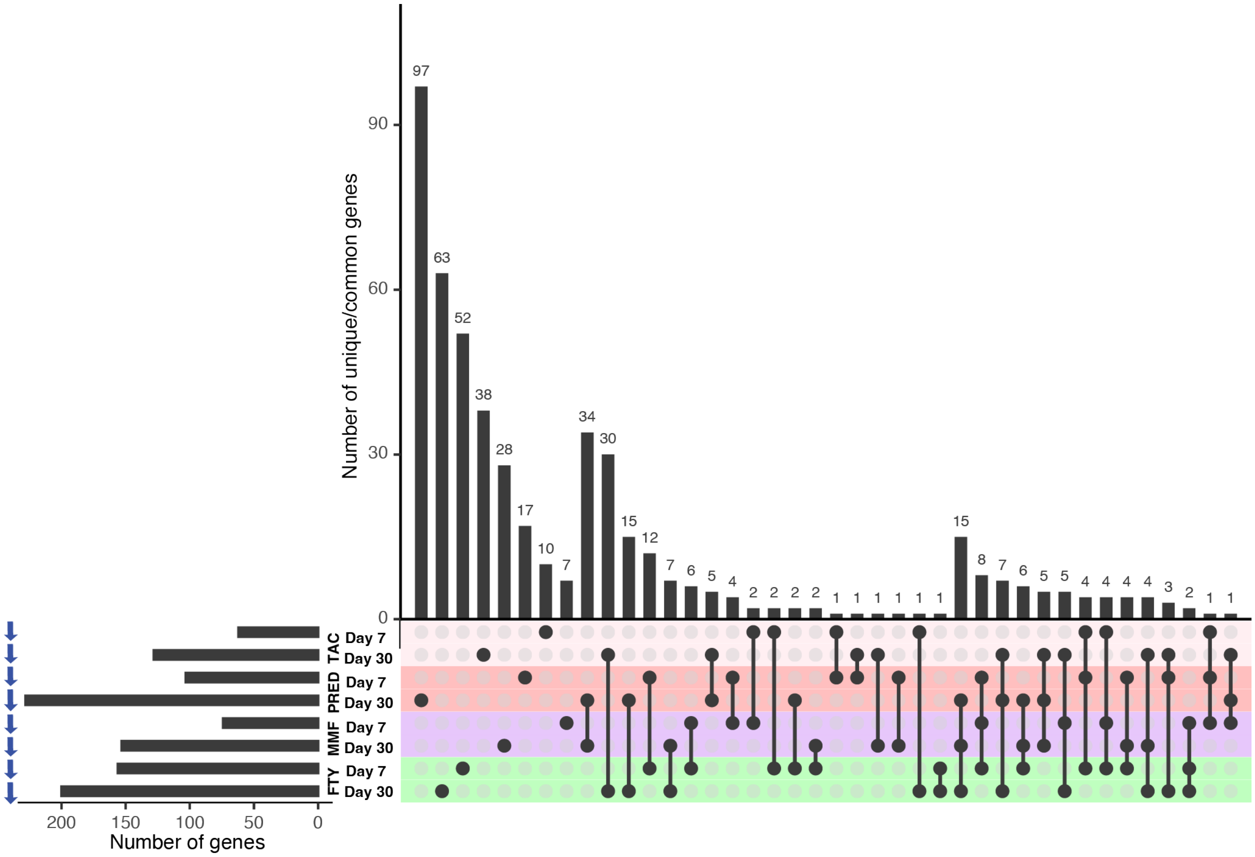

### Supplemental Figure 3. C)

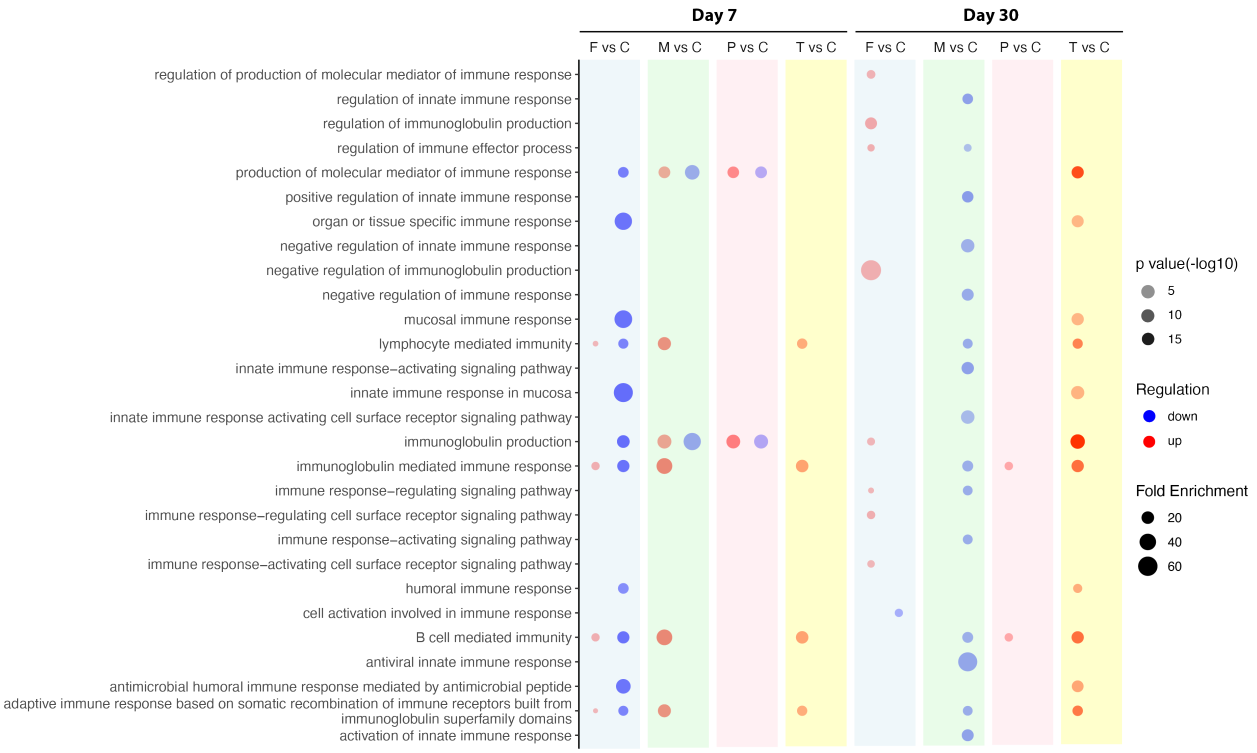

**Supplemental Figure 4.**

A

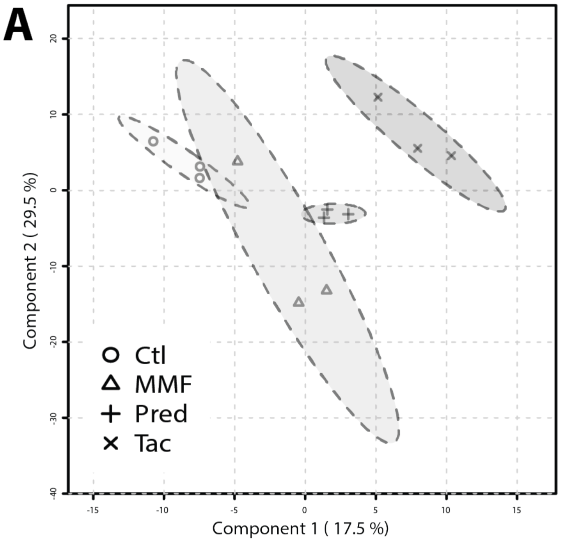

#### Supplemental Figure 4.

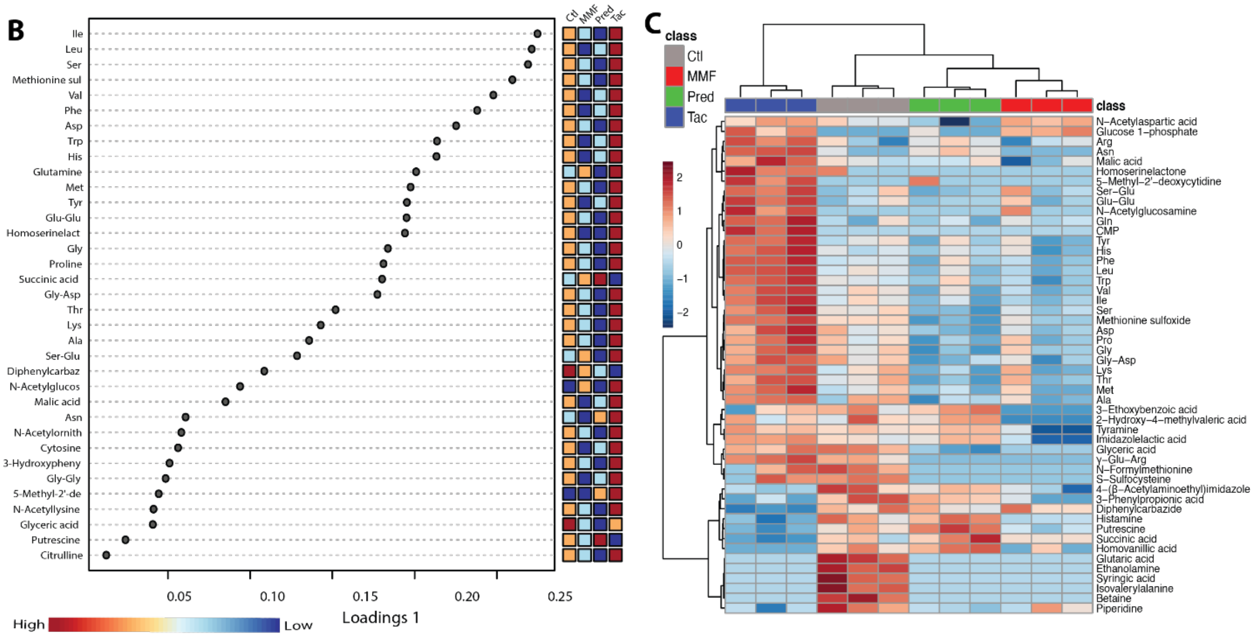

### Supplemental Figure 5.

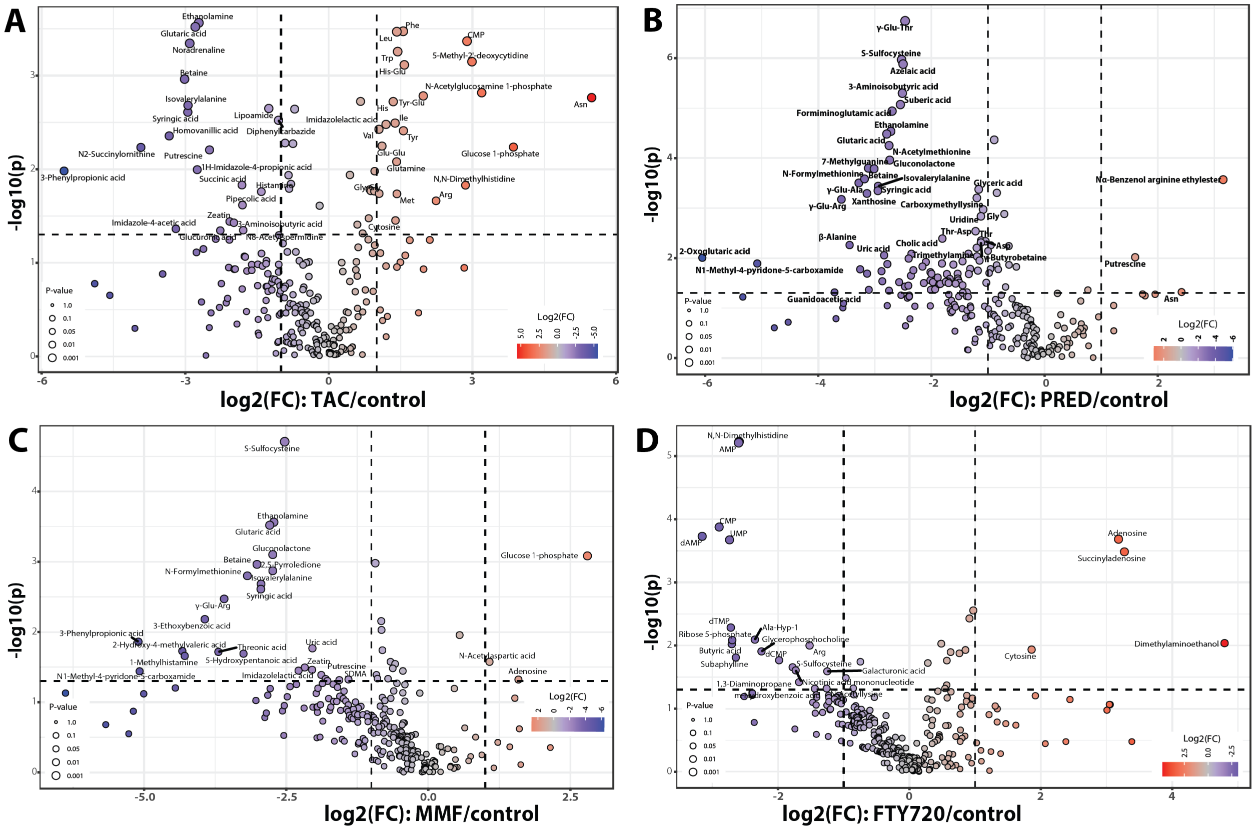

**Supplemental Figure 6.**

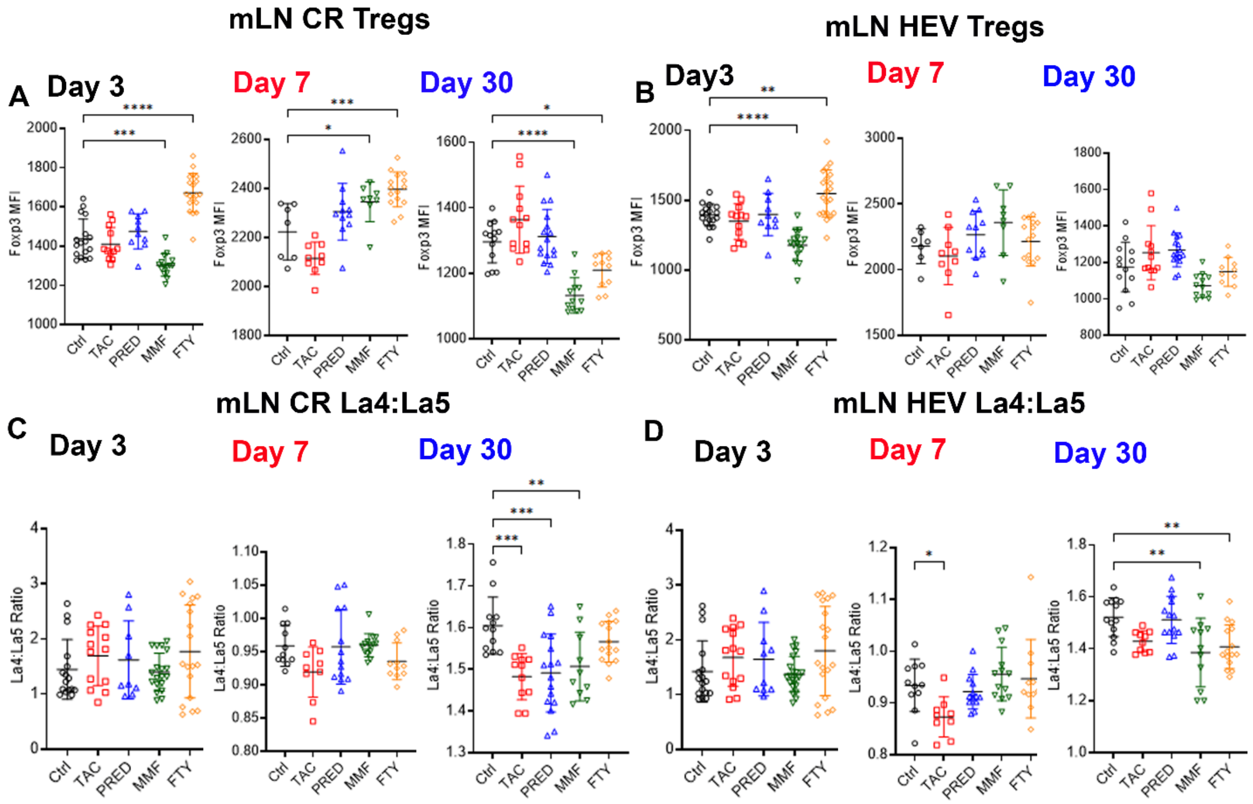

**Supplemental Figure 7.**

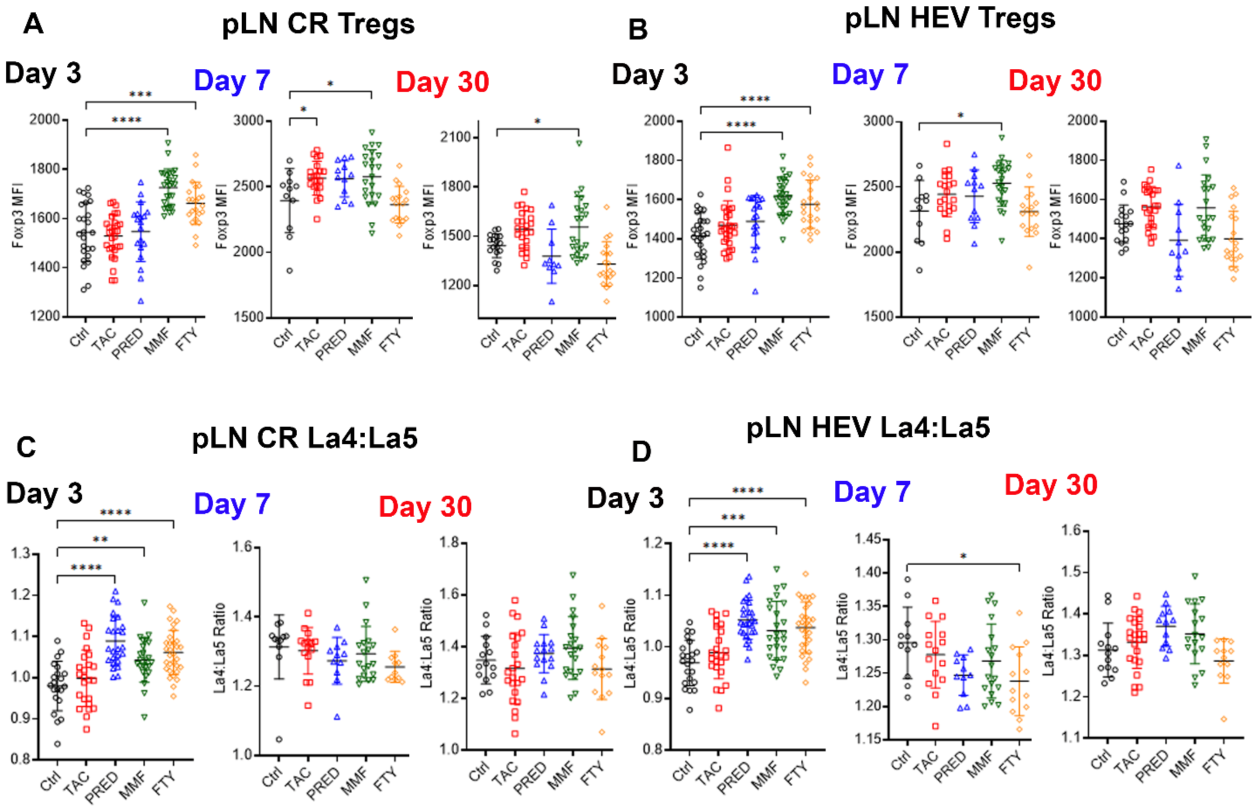

#### Supplemental tables

**Supplemental Table 1. Gut microbiome characteristics. A**) Gut microbiota taxonomy on days 3, 7, and 30, characterized using the comprehensive mouse microbiota genome catalog [6]. **B**) Functional characterization performed using HUMAnN3 (Human Microbiome Project Unified Metabolic Analysis Network) (v3.0.0.alpha.3) [4] and Uniref90 database[5] for prevalence and abundance of metabolic pathways.

**Supplemental Table 2. Transcriptome of intestinal tissue. A**) Statistics of the transcriptome data. Detailed list of differentially expressed genes (DEGs) of via comparison of individual treatment group versus no treatment control that are upregulated at **B**) day 7 and **C**) day 30, downregulated at **D**) day 7 and **E**) day 30. DEGs between control and treatment groups at day 7 and day 30. Significant differential expression assessed using DEseq2 (5) with an FDR value ≤ 0.05 and Fold-of-change (FC)>2. **Abbr**: TAC: tacrolimus, MMF: mycophenolate mofetil, PRED: prednisone, FTY: fingolimod.

**Supplemental Table 3. Metabolome of gut lumen and serum. A**) Luminal metabolome of the intraluminal stool for TAC, MMF, PRED, and a no-treatment control group. **B**) Metabolome of stool pellets after FTY treatment, specimen collected at the same time points for metabolomic analysis due to technical limitations in obtaining sufficient sample volumes for comprehensive profiling. Significantly altered metabolites using pairwise comparison of individual drugs to the no-treatment control for **C**) FTY; **D**) PRED, **E**) MMF, and **F**) TAC. Fold change >2 and the p-value <0.05 based using 2-sample t-tests.

**Supplemental Table 4. List of Antibodies**

| Target Molecule | Clone | Catalog No. |
|---|---|---|
| Anti-Hamster IgG Cy3 | Polyclonal | Jackson ImmunoResearch; 127-165-160 |
| Anti-Mouse IgG-AF647 | Polyclonal | Jackson ImmunoResearch; 715-605-151 |
| Anti-Rabbit AF488 | Polyclonal | Jackson ImmunoResearch; 111-545-003 |
| Anti-Rabbit Cy5 | Polyclonal | Jackson ImmunoResearch; 111-175-003 |
| Anti-Rabbit DL405 | Polyclonal | Jackson ImmunoResearch; 711-476-152 |
| Anti-Rabbit IgG-AF488 | Polyclonal | Jackson ImmunoResearch; 711-545-152 |
| Anti-Rabbit IgG-AF594 | Polyclonal | Jackson ImmunoResearch; 111-585-003 |
| Anti-Rabbit IgG-AF647 | Polyclonal | Jackson ImmunoResearch; 711-606-152 |
| Anti-Rat IgG AF594 | Polyclonal | Jackson ImmunoResearch; 112-586-143 |
| Anti-rat IgG AF647 | Polyclonal | Jackson ImmunoResearch; 712-606-153 |
| CD4 | GK1.5 | Biolegend; 100401 |
| B220 | RA3-6B2 | eBioSci; 17-0452-82 |
| CD11b | M1/70 | eBioSci; 11-0112-81, 17-0112-82 |
| CD8 | 53-6.7 | Biolegend; 100701 |
| CD11c | HL3 | BD; 550283, 557401 |
| ER-TR7 | ER-TR7 | Novus; NB100-64932 |
| ER-TR7 | Polyclonal | Santa Cruz; SC-73355 |
| Foxp3 | NRRF-30 | eBioSci; 14477180 |
| Foxp3 | FJK-16s | eBioSci; 12-5773-82 |

| | | |
|---|---|---|
| Laminin $\alpha$4 | 775830 | R&D; MAB3837 |
| Laminin $\alpha$5 | Polyclonal | Novus Biol; NBP1-18714 |
